# Cortical circuitry mediating inter-areal touch signal amplification

**DOI:** 10.1101/2023.06.06.543886

**Authors:** Lauren Ryan, Andrew Sun-Yan, Maya Laughton, Simon Peron

## Abstract

Sensory cortical areas are often organized into topographic maps which represent the sensory epithelium^1,2^. Individual areas are richly interconnected^3^, in many cases via reciprocal projections that respect the topography of the underlying map^4,5^. Because topographically matched cortical patches process the same stimulus, their interaction is likely central to many neural computations^6-10^. Here, we ask how topographically matched subregions of primary and secondary vibrissal somatosensory cortices (vS1 and vS2) interact during whisker touch. In the mouse, whisker touch-responsive neurons are topographically organized in both vS1 and vS2. Both areas receive thalamic touch input and are topographically interconnected^4^. Volumetric calcium imaging in mice actively palpating an object with two whiskers revealed a sparse population of highly active, broadly tuned touch neurons responsive to both whiskers. These neurons were especially pronounced in superficial layer 2 in both areas. Despite their rarity, these neurons served as the main conduits of touch-evoked activity between vS1 and vS2 and exhibited elevated synchrony. Focal lesions of the whisker touch-responsive region in vS1 or vS2 degraded touch responses in the unlesioned area, with whisker-specific vS1 lesions degrading whisker-specific vS2 touch responses. Thus, a sparse and superficial population of broadly tuned touch neurons recurrently amplifies touch responses across vS1 and vS2.

## Introduction

In the canonical columnar microcircuit^11^, thalamic input strongly activates cortical layer (L) 4^12,13^, which drives activity in L2/3^14^. Within L2/3, similarly tuned neurons exhibit elevated recurrent connectivity^15^, which enables pattern completion^16^ and amplification^17,18^. L2/3 of many cortical areas both outputs extensively to^19,20^ and receives input from^21^ other cortical areas, particularly adjacent ones^22^. It is thus likely that local recurrent processing in L2/3 is complemented by interareal recurrence. Topographically matched recurrent amplification between cortical areas could facilitate coordination across patches of cortex responding to the same stimulus. Such recurrence could drive location specific response enhancement characteristic of spatial attention^6,7^, undergird feature binding across areas^6,8^ and, by ensuring fairly synchronous activation of cortical patches responding to the same stimulus across distinct cortical areas, facilitate object recognition^9,10^.

In the mouse vibrissal system, thalamic input from individual whiskers terminates on a small patch of vS1 cortex called a ‘barrel’^23^. Vibrissal S2 also shows somatotopically organized responses to whisker touch, but with a more compressed map^24,25^. These areas are extensively interconnected in a topography preserving manner^4^ and respond robustly to whisker touch^25-27^. Axonal projections both from vS1 to vS2 and from vS2 to vS1 carry a strong touch signal^4,21,24,28^. This suggests that vS1 and vS2 may recurrently enhance one another’s touch responses^24^.

How do somatotopically matched (‘iso-somatotopic’) subregions of vS1 and vS2 interact during active touch? Here, we focus on subregions of vS1 and vS2 that respond to touch from whiskers C2 and C3. We first employ volumetric two-photon calcium imaging^29,30^ of iso-somatotopic subregions of vS1 and vS2 to characterize touch neuron populations in both areas. Next, we use retrograde labeling to determine which populations of touch neurons relay touch information to iso-somatotopic targets across both areas. Finally, we selectively lesion patches of either vS1 or vS2 responsive to touch by the spared whiskers to assess how iso-somatotopic sites mutually influence one another.

### Mapping neural activity in vS1 and vS2 during an active touch task

We implanted transgenic mice expressing GCaMP6s^31^ in cortical excitatory neurons (Ai162 X Slc17a7-Cre)^32^ with a cranial window over vS1 and vS2 (Methods). Following recovery, mice were trimmed to two whiskers (C2 and C3) and trained on a head-fixed active touch task. Trials started with a 1 second stimulus period during which a pole became accessible to palpation near the tip of one of the two spared whiskers (**Fig. 1a**). The pole was then withdrawn, and a 2 second delay period began. Finally, a tone indicated the start of a 1 second response period, during which licking resulted in a water reward on a random 70% of trials. This task design ensured that animals whisked actively and were reward motivated, both of which influence sensory cortical responses^33,34^.

**Figure 1.**
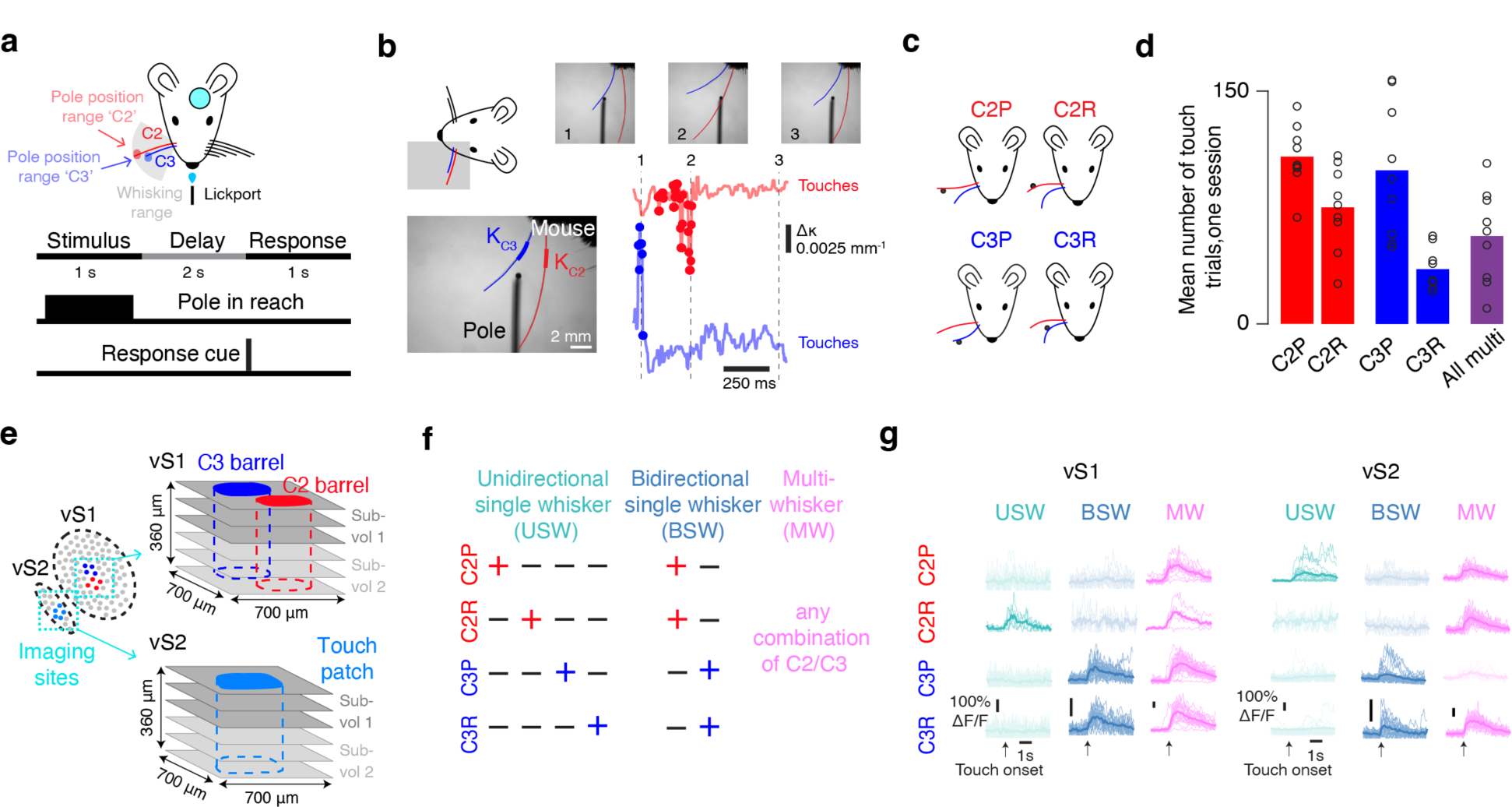
Volumetric two-photon imaging in vS1 and vS2 during an active two-whisker touch task. **a,** Experimental setup. Head-fixed mice palpated poles that were presented within range of one of their two spared whiskers. Mice received reward on a random 70% of trials. Bottom, task timing. A pole was presented for 1 second and was then removed. Following a 2 second delay which ended with a response cue, mice had 1 second to respond by licking. **b,** Whisker videography. Bottom left, example frame showing traced whiskers and where curvature of each whisker is calculated. Top right, three example frames that correspond to points in time on the Δκ traces below. Bottom right, Δκ trace for each whisker, with touches overlaid. **c,** Four types of single-whisker touch: whisker C2 protraction (C2P), whisker C2 retraction (C2R), whisker C3 protraction (C3P) and whisker C3 retraction (C3R). **d,** Number of trials of a given touch type over one imaging session, n=9 mice. Bars, mean; circles, individual mice. Red, whisker C2 touch trials; blue, whisker C3 touch trials; purple, dual-whisker touch trials. **e,** Volumetric calcium imaging. Left, placement of cranial window allows for imaging both vS1 and vS2. Subregions responsive to spared whiskers colored. Right, groups of three planes of the same shade constitute a simultaneously imaged subvolume. **f,** Touch cell type classification. Each row represents an example cell of the shown category, with plus indicating that the neuron responds to that touch type. **g,** Example ΔF/F traces for each cell type from each area (left, vS1; right, vS2) for one example session from one animal. Thin lines, individual trial responses; thick line, mean across trials.

We employed high-speed whisker videography to capture whisker movement (**Fig. 1b**; 400 Hz; Methods). Whisker video was segmented to track individual whiskers^35^ (Methods), allowing us to measure changes in whisker curvature (Δκ) which was used as a proxy for force at the whisker follicle^36^. We divided touches into four types (**Fig. 1c**): protractions of whisker C2 (C2P), protractions of whisker C3 (C3P), retractions of whisker C2 (C2R), and retractions of whisker C3 (C3R). We used a range of pole positions to ensure many isolated touches of each type (**Fig. 1a, d**), with most trials containing touch (79.4% ± 4.4% trials with touch, mean ± S.D., n=9 mice).

In trained mice (n=9; **Extended Data Table 1**) with consistent whisking, we recorded activity during the task using volumetric two-photon calcium imaging^30^ (**Fig. 1e**). We first determined the precise locations in vS1 and vS2 that responded to touch by the two spared whiskers by passively deflecting the whiskers during widefield two-photon calcium imaging (Methods; **Extended Data Fig. 1**). In vS1 we found the locations corresponding to the C2 and C3 barrels. In vS2, we located the patch of touch cells that responded to touch from either C2, C3, or both. In both areas, we performed cellular resolution recordings centered on these touch responsive subregions, simultaneously imaging three individual planes (700-by-700 μm) spaced 60 μm apart in depth (7 Hz). These planes comprised a ‘subvolume’, and we imaged two subvolumes per area (**Extended Data Table 1**), starting at the L1-L2 boundary. While all subvolumes were visited in one session, only one subvolume was imaged at a time (Methods). We recorded across all or most of L2/3 depth-wise in both areas (8,864 ± 563 neurons per mouse, n=9 mice).

We classified neurons as responsive to a particular touch type if they responded on at least 5% of trials that contained only that touch type (Methods). Neurons that responded to at least one touch type were considered touch neurons and were further classified based on the specific touch type(s) they responded to. Neurons that responded to only one type of whisker touch were classified as unidirectional single whisker cells (USW); neurons that responded to both protraction and retraction touches for only one whisker were classified as bidirectional single whisker cells (BSW); neurons that responded to any combination of C2 and C3 touches were classified as multiwhisker cells (MW) (**Fig. 1f**). We found neurons of each type in both vS1 and vS2 (**Fig. 1g**).

### Multiwhisker cells are rare but respond robustly to touch in both vS1 and vS2

In superficial vS1, broader tuning and sparser responses are especially pronounced in L2^37,38^. To determine if a similar organization was present in vS2, we examined the touch neuron distribution in vS1 and vS2, finding a mix of all three types in both areas (**Fig. 2a, b**). We imaged vS1 and vS2 in 9 mice across a depth of 360 μm starting at the L1-L2 boundary. Because the vS2 topographic map is compressed in comparison to the map in vS1, we sub-selected an approximately 300-by-600 μm area centered on the most touch responsive neurons, yielding 1,963 ± 408 vS2 neurons (Methods). This subregion contained the majority of C2 and C3 responsive neurons. For vS1, we used a 600-by-600 μm area centered on the barrels of the two spared whiskers (neuron count: 4,017 ± 607).

**Figure 2.**
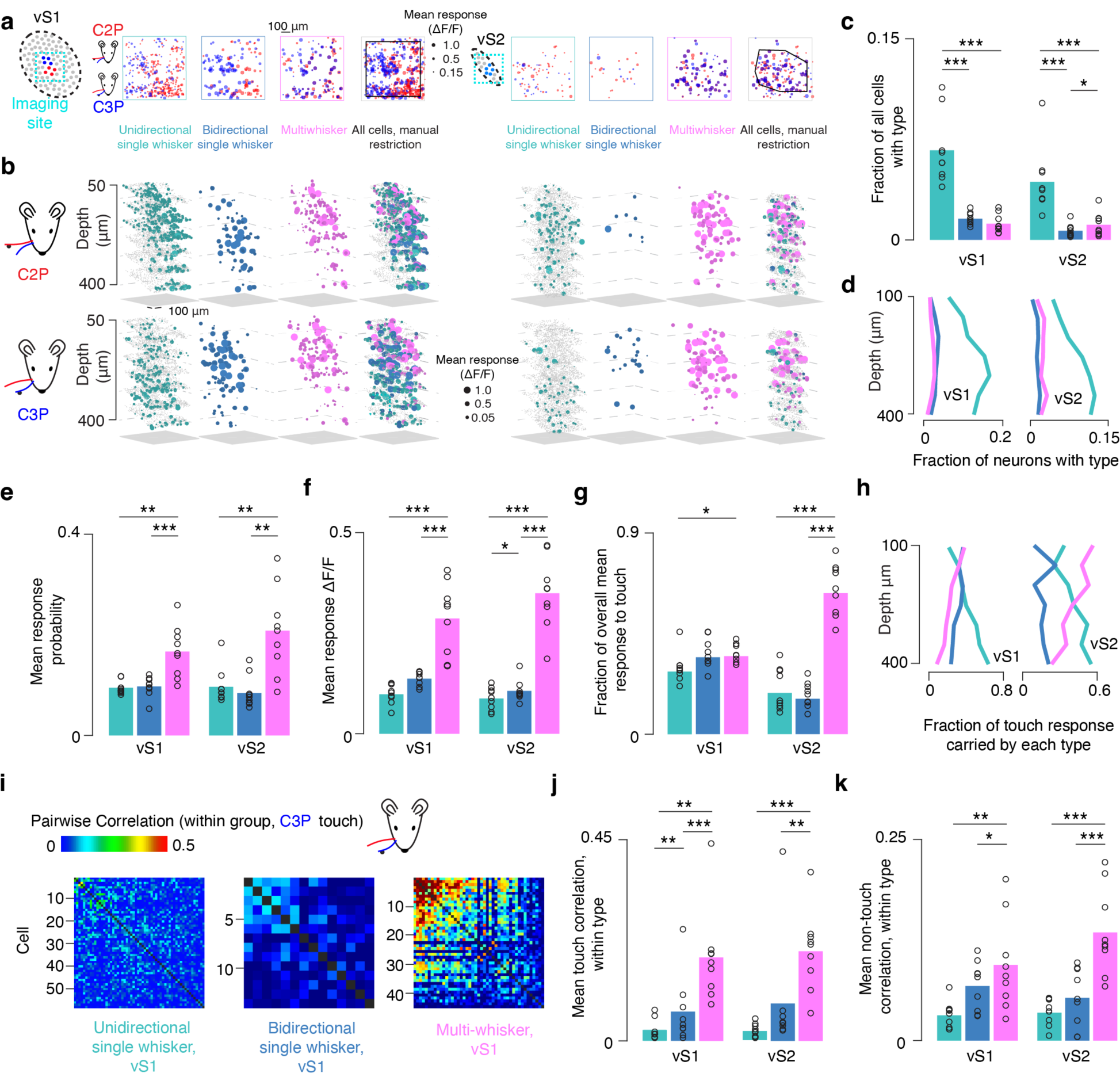
A sparse population of multiwhisker cells exhibits robust touch responses in both vS1 and vS2. **a,** Projection across depth of mean touch-evoked ΔF/F among to C2P (red) and C3P (blue) touches in an example. Only cells with mean touch-evoked ΔF/F greater than 0.15 are included. Left to right: unidirectional single-whisker neurons; bidirectional single-whisker neurons; multiwhisker neurons; all neurons, with restriction border shown in black (Methods). Left four panels, vS1; right four panels, vS2. **b,** Example neurons from **a** shown in 3D. Top, mean touch-evoked ΔF/F following C2P touch; bottom, C3P touch. Left to right, mean touch-evoked ΔF/F among unidirectional single-whisker neurons (cyan; grey dots, unresponsive neurons), bidirectional single-whisker neurons (dark blue), multiwhisker neurons (magenta), all neurons in sub selected region. Left four panels, vS1; right four panels, vS2. **c,** Frequency of each of the major touch neuron types in both areas. Bars, mean (n=9 mice). P-values indicated for two-tailed paired t-tests: * p<0.05, ** p<0.01, *** p<0.001. **d,** Fraction of neurons belonging to type at given depth (bin size: 50 μm) in each area. **e,** Mean response probability for each type in each area averaged across all touch types for which individual neurons were deemed responsive. **f,** Mean touch-evoked ΔF/F averaged across touch types for which individual neurons were deemed responsive. **g,** Fraction of overall mean ΔF/F response to touch contributed by each population. **h,** Fraction of touch response carried by each type at each depth (bin size: 50 μm). **i,** Example within-type C3P touch-evoked correlation matrices for each touch cell type in vS1 in an example animal (Methods). **j,** Mean within-type pairwise correlations in each area for periods of touch. **k,** Same as in **j**, but ‘spontaneous’ correlations for periods of non-touch.

Vibrissal S2 contained a smaller fraction of unidirectional single whisker (**Fig. 2c**; USW fraction, vS1: 0.067 ± 0.026, vS2: 0.044 ± 0.025, n=9 mice, p=0.008, paired t-test comparing vS1 and vS2) and bidirectional single whisker cells (BSW fraction, vS1: 0.016 ± 0.005, vS2: 0.007 ± 0.005, p=0.004) than vS1, with a comparable number of multiwhisker neurons (MW fraction, vS1: 0.012 ± 0.007, vS2: 0.011 ± 0.009, p=0.820). Both areas had substantially more unidirectional single whisker neurons than bidirectional or multiwhisker neurons (vS1, USW vs. BSW, p<0.001; USW vs. MW, p<0.001; vS2, USW vs. BSW, p<0.001; USW vs. MW, p<0.001). In vS2, multiwhisker cells were more frequent than bidirectional single whisker cells (BSW vs. MW, p=0.015); in vS1, this was reversed, though the difference was not significant (p=0.931). Broadly tuned neurons (i.e., multiwhisker and bidirectional neurons) thus made up a small fraction of neurons in both areas.

We next determined the depth distribution of the different touch populations starting at the L1-L2 border. All three touch neuron types declined in frequency from deep L3 to superficial L2, with broadly tuned neurons declining more slowly (**Fig. 2d**). We restricted our analyses to the most superficial three imaging planes where these broadly tuned neurons were relatively more numerous and asked which population contributed the most to the touch response. In both areas, the multiwhisker neurons had a significantly higher touch response probability compared to bidirectional (**Fig. 2e**, n=9, vS1: p<0.001, paired t-test; vS2: p=0.002) and unidirectional single whisker cells (vS1: p=0.003; vS2: p=0.003). Multiwhisker neurons in both areas also had a significantly larger mean touch-evoked ΔF/F response (averaged only across touch types to which a neuron was considered responsive; Methods) than bidirectional (**Fig. 2f**, n=9 mice, vS1: p<0.001, vS2: p<0.001) and unidirectional single whisker cells (vS1: p<0.001, vS2: p<0.001). Multiwhisker neurons therefore exhibit the strongest touch responses in superficial L2/3. These trends persisted when including neurons from all depths (**Extended Data Fig. 2**).

Could the high responsiveness of multiwhisker neurons compensate for their rarity, resulting in this population carrying much of the touch response? To address this, we computed the mean touch-evoked ΔF/F for every touch neuron for each individual touch. We then summed across all neurons of a given touch type and divided each group’s net response by the net response across all neurons, resulting in an estimate of the fractional contribution each population made for every touch. We then computed the mean contribution of each population across all touches (Methods). In vS1, we found that each group carries a relatively comparable proportion of the touch response (**Fig. 2g**). In vS2, however, multiwhisker neurons were responsible for most of the overall response to touch (MW vs. USW: p<0.001, MW vs. BSW: p<0.001). Therefore, despite their rarity, multiwhisker neurons carry a disproportionately large fraction of the touch response in superficial L2/3 in both areas and comprise the majority of the vS2 touch response.

We next asked what fraction of the touch response is carried by each population across depth. Broadly tuned neurons contribute an increasing fraction of the touch response superficially in both areas, with the response of multiwhisker neurons becoming especially dominant in superficial vS2 (**Fig. 2h**). In contrast, unidirectional single whisker cells showed a declining fractional response in both areas superficially. As input from L4 is transformed by local circuitry on its way to L2^39^, the touch response thus becomes increasingly concentrated among multiwhisker neurons in both areas.

If multiwhisker neurons exhibit elevated synchrony relative to other touch populations, they would be more effective at driving putative downstream targets^40^. We therefore compared touch-evoked correlations between neurons within each population (**Fig. 2i**; Methods). Multiwhisker cells exhibited significantly higher touch-evoked correlations than other populations (**Fig. 2j**; n=9 mice, vS1: MW vs. USW p=0.003, MW vs. BSW p<0.001, vS2: MW vs. USW p=<0.001, MW vs. BSW p=0.003).This pattern was also present in ‘spontaneous correlations’ measured during the non-touch epoch (Methods): multiwhisker cells were significantly more correlated to one another compared with other touch cell types (**Fig. 2k**, n=9 mice, vS1: MW vs. USW p=0.008, MW vs. BSW p=0.043, vS2: MW vs. USW p<0.001, MW vs. BSW p<0.001). Thus, in addition to their larger touch responses, multiwhisker cells exhibit greater synchrony, making them especially well-suited to influence downstream populations. Elevated spontaneous correlations suggest that this synchrony may be due to elevated connectivity among these neurons^15^.

In sum, despite multiwhisker cells constituting a minority of touch responsive neurons, these cells responded to touch strongly, reliably, and with elevated synchrony in both vS1 and vS2. These neurons thus play a disproportionate role in the superficial touch response in both areas.

### Multiwhisker neurons contribute disproportionately to the vS1-vS2 projection touch response

Vibrissal S1 and vS2 are richly interconnected^4,24,41,42^ and respond robustly to whisker touch^4,25,43^. Do broadly tuned touch neurons contribute disproportionately to the touch signal carried between iso-somatotopic vS1 and vS2 patches? We labelled projecting neurons in both areas by focally injecting rAAV2-retro-FLEX-tdTomato into L2/3 of either vS1 or vS2 (**Fig. 3a, b**). We targeted either the center of the spared barrels in vS1 or the center of the ‘touch patch’ responding to the spared whiskers in vS2 (Methods). Injections into the vS2 touch patch led to a diffuse labeling across vS1 (**Fig. 3a**; **Extended Data Fig. 3**), implying that vS1 sends broad touch input to vS2. In contrast, vS1 injections led to focal expression in the vS2 touch patch (**Fig. 3b**).

**Figure 3.**
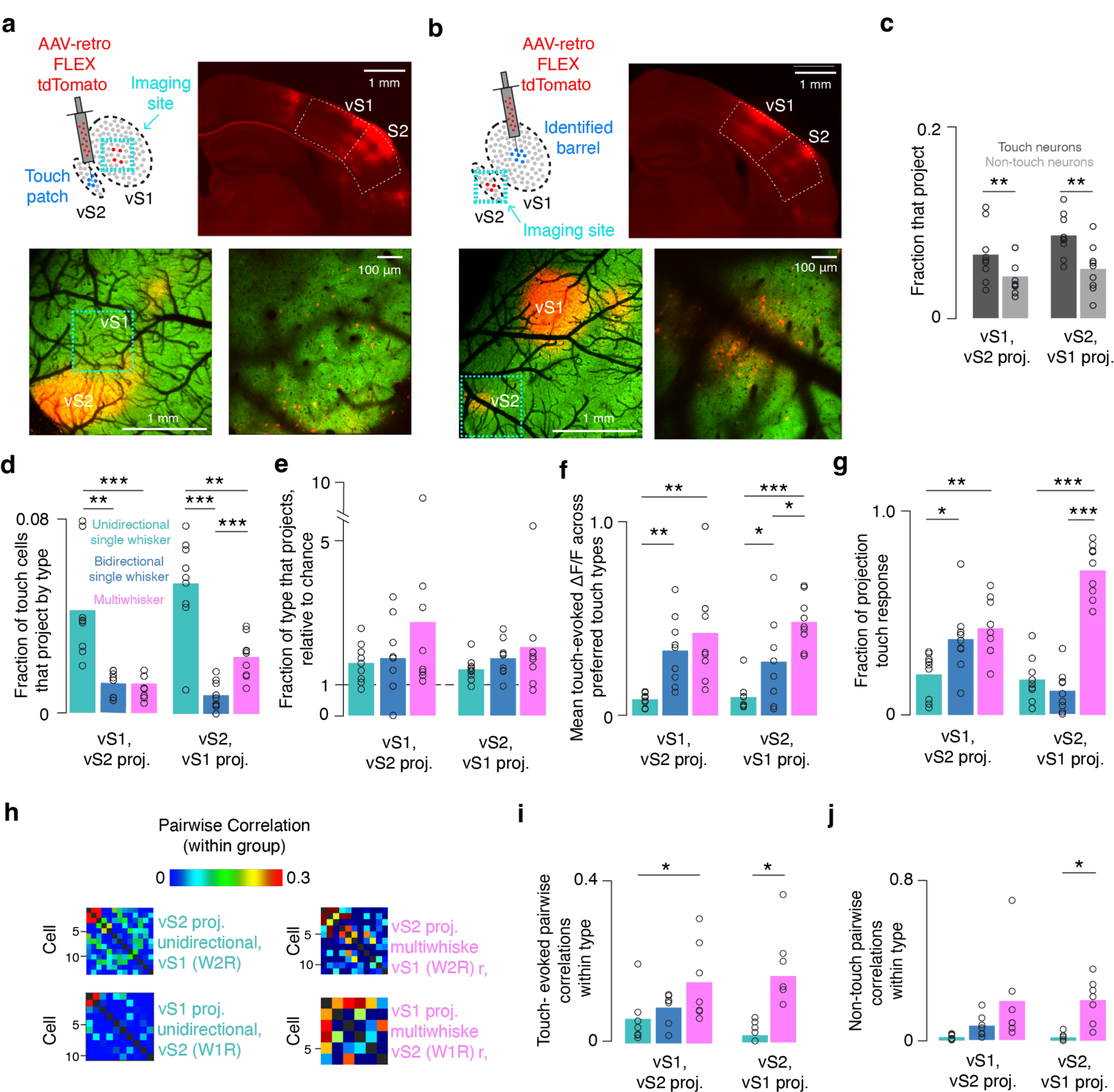
Multiwhisker cells carry a large fraction of the vS1-vS2 projection touch response. **a,** Retrograde injection into vS2. Top-right, coronal section showing injection in vS2 and retrograde viral tdTomato expression in vS1. Bottom-left, widefield 2-photon image showing labeled cell expression. Green, GCaMP6s; red, tdTomato. Cellular resolution imaging site is outlined in cyan. Bottom-right, example cellular resolution two-photon imaging plane in vS1 showing cells expressing GCamp6s and tdTomato. **b,** As in **a**, but for vS2 imaging following vS1 injection. Injection is made into a touch patch identified by cellular-resolution imaging prior to injection (Methods). **c,** Fraction of touch (dark grey) and non-touch (light grey) neurons that project from each area to the other. Bars, mean (n=9 mice); circles, individual animals. P-values indicated for two-tailed paired t-tests: * p<0.05, ** p<0.01, *** p<0.001. **d,** Fraction of touch cells of given type that project. **e,** Fraction each of each touch subpopulation that projects to the other area, normalized to fraction of all neurons belonging to that subpopulation (‘chance’). **f,** Mean ΔF/F in response to touch averaged across touch types for which individual projecting neurons are responsive. **g,** Fraction of touch-evoked ΔF/F (Methods) among projecting neurons by category for both areas. **h,** Example within-type C3R touch correlation matrix for two example populations in one mouse that project from vS1 to vS2 and C2R responses for analogous populations in a second mouse projecting from vS2 to vS1. **i,** Mean within-type pairwise correlations during the period around touch (n=7 mice; Methods). **j,** Mean within-type pairwise correlations during non-touch ‘spontaneous’ period for each touch population in each area (Methods).

We imaged the subregion of the un-injected area that responded to touch by the two spared whiskers. In both areas, we imaged two 3-plane subvolumes spanning 90 to 120 μm each (total: 180 to 240 μm in depth; **Extended Data Table 1**), restricting analysis to an approximately 600-by-600 μm patch in vS1 and an approximately 300-by-600 μm patch in vS2 (Methods). We found 180 ± 49 vS2-projecting neurons in vS1 out of a total 4,109 ± 788 neurons (n=9 mice), and 97 ± 58 vS1-projecting neurons in vS2 out of 1,705 ± 550 (n=9 mice) neurons. In both areas, touch neurons were more likely to project to the other area than non-touch neurons (**Fig. 3c**): in vS1, 7.0% ± 3.0% of touch neurons projected to vS2, whereas only 4.0% ± 2.0% of non-touch neurons projected to vS2 (paired t-test, touch v. non-touch p=0.003). In vS2, 9.0% ± 2.0% of touch-neurons and 5.0 ± 2.0% of non-touch neurons projected to vS1 (p=0.002). Thus, touch neurons are more likely to project in both directions than non-touch neurons.

We next examined the composition of the projecting populations. Among vS2-projecting neurons in vS1, unidirectional single whisker neurons were more numerous than bidirectional and multiwhisker neurons (**Fig. 3d**; two-sample t-test, n=9 mice, USW vs. BSW p=0.001; USW vs. MW p=0.001). A similar pattern held in vS2 (two-sample t-test, n=9 mice, USW vs. BSW p<0.001; USW vs. MW p=0.001). In vS2, multiwhisker neurons were more likely to project than bidirectional single whisker neurons (p<0.001). Given that unidirectional single whisker neurons were more numerous in both areas (**Fig. 2c**), we asked if specific touch cell types projected more than expected by chance. We divided the fraction of the projecting population consisting of a particular type by the fraction of the overall population consisting of that type. This number exceeded 1 for all touch types (**Fig. 3e**), implying that all touch neuron types projected in both directions more than predicted by their frequency.

Did touch-evoked response amplitude differ across specific projecting touch populations? Among vS1 neurons projecting to vS2, multiwhisker cells responded more strongly on average than unidirectional single whisker cells (**Fig. 3f**, n=9 mice, USW vs. MW p=0.001, t-test), as did bidirectional single whisker cells (USW vs. BSW, p=0.001). Among vS2 neurons projecting to vS1, multiwhisker cells again responded more strongly on average than unidirectional single whisker cells (n=9 mice, USW vs. MW p<0.001, t-test), as did bidirectional single whisker cells (USW vs. BSW, p=0.040). Multiwhisker projecting neurons also responded more strongly on average compared to bidirectional single whisker neurons in vS2 (BSW vs. MW, p=0.037). Multiwhisker neurons thus exhibit the largest touch responses among projecting neurons, particularly in vS2.

What fraction of touch-evoked activity among projecting neurons did specific populations contribute? For vS1 neurons projecting to vS2, multiwhisker neurons carried the largest fraction of the touch response (**Fig. 3g**, 0.43 ± 0.14, n=9 mice), significantly more than the fraction carried by unidirectional single-whisker neurons (0.20 ± 0.11, p=0.002, t-test comparing MW vs. USW fraction). Bidirectional single-whisker neurons also carried a larger fraction of the touch response than unidirectional neurons (0.37 ± 0.17, p=0.020, t-test comparing USW vs. BSW fraction). Among vS2 neurons projecting to vS1, multiwhisker neurons carried the largest fraction of the touch response (MW: 0.71 ± 0.13; BSW: 0.11 ± 0.10; USW: 0.17 ± 0.11; MW vs. USW, p<0.001; MW vs. BSW, p<0.001). Despite their rarity, multiwhisker neurons carry the largest fraction of the touch response for both projections.

As broadly tuned touch neurons exhibited greater synchrony (**Fig. 2j**), making them potentially more effective at influencing downstream activity^40^, we next asked if this trend held for projecting neurons. We examined touch-evoked correlations in specific populations of projecting neurons (Methods; only some animals had enough projecting neurons to calculate correlations). Among vS1 neurons projecting to vS2, multiwhisker projecting neurons had higher within-group pairwise correlations than unidirectional single whisker neurons near the time of touch (**Fig. 3h-i**, MW vs. USW, p=0.021, paired t-test, n=7). Due to a paucity of projecting bidirectional single whisker cells in vS2, we only examined unidirectional single whisker and multiwhisker neurons in vS2, with multiwhisker neurons exhibiting higher correlations around the time of touch (MW vs. USW, p=0.046, n=7). We found no significant differences in the touch-evoked pairwise correlations between projecting and non-projecting neurons near the time of touch. In both vS1 and vS2, multiwhisker projecting neurons had higher spontaneous correlations outside the touch epoch, though only the difference between multiwhisker and unidirectional single whisker cells was significant (**Fig. 3j**; MW vs. USW, p=0.025, n=7).

Projecting multiwhisker neurons are thus highly correlated both during touch and non-touch epochs, and so more likely to send a synchronized signal downstream.

Overall, broadly tuned neurons make up a disproportionate fraction of projecting neurons across iso-somatotopic sites in vS1 and vS2. Projecting multiwhisker cells exhibited elevated synchrony, suggesting that they evoke downstream activity more effectively.

### Vibrissal S2 lesions degrade vS1 touch responses

Given that vS2 touch activity among multiwhisker neurons contributes disproportionately to the vS1-projecting population, we next asked how vS2 contributes to iso-somatotopic touch responses in vS1. We ablated the vS2 touch patch using prolonged exposure to a femtosecond laser source^44^ (Methods), resulting in a focal, superficial vS2 lesion (**Fig. 4a-c**). Before and after vS2 lesions, we imaged vS1 using a single 3-plane subvolume spanning 180 μm in total depth (**Extended Data Table 1**). We examined the impact of vS2 lesions on touch responses in 6 mice across 2,477 ± 465 vS1 neurons confined to a ∼600-by-600 μm subregion (**Fig. 4d**). Lesions significantly reduced the relative mean touch-evoked ΔF/F in vS1 (**Fig. 4e**; p=0.009, t-test comparing relative change to 0). We found the same relative decline in touch-evoked ΔF/F among all three touch cell types (**Fig. 4f**; USW, p=0.013; BSW, p=0.031; MW, p=0.011). Vibrissal S2 thus augments the vS1 response to touch by providing broad touch input to all touch cell types.

**Figure 4.**
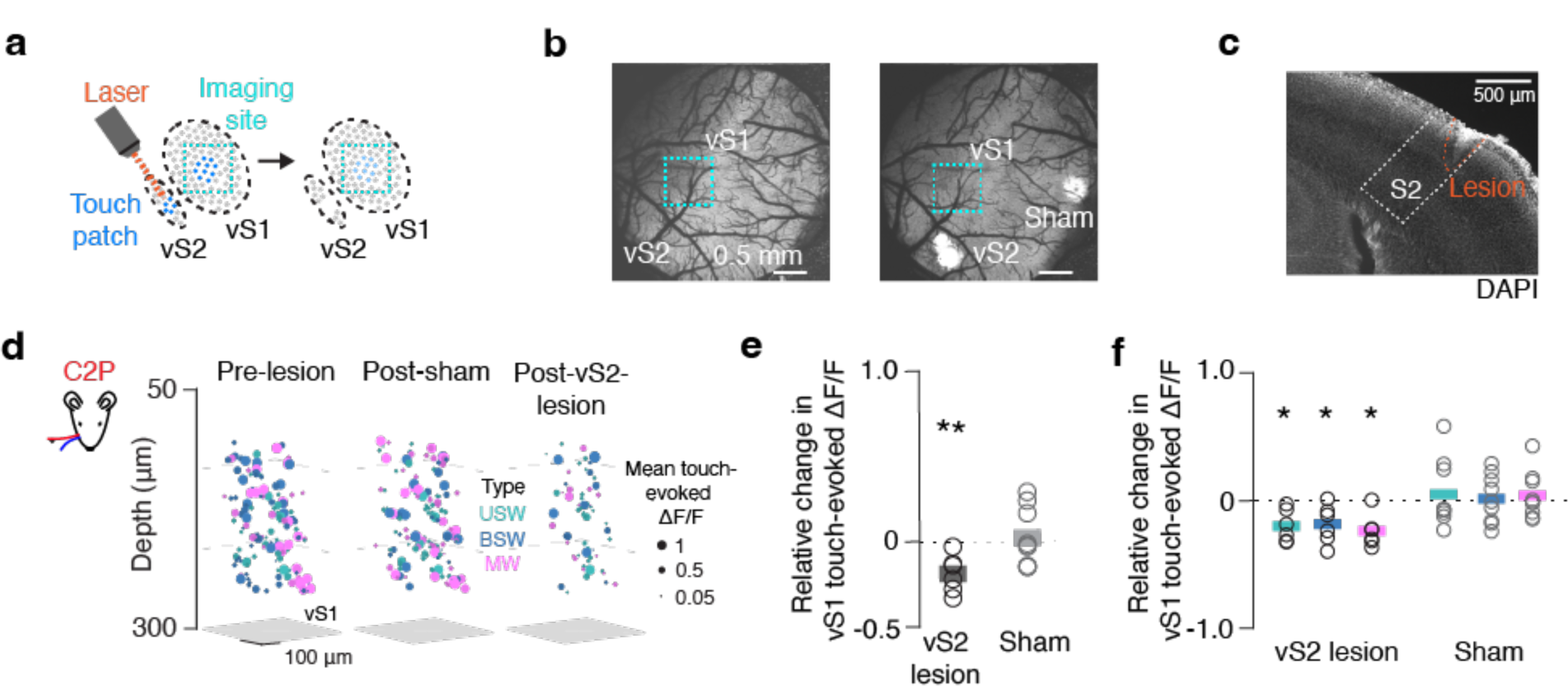
Touch response in vS1 declines after vS2 lesion. **a,** >Lesion targeting the touch patch in vS2. **b,** Widefield 2-photon image (4X, Methods) depicting the window before and after lesions in an example animal. Vibrissal S2 lesion took place 24 hours prior to right image; sham lesion took place 72 hours prior to vS2 lesion. **c,** Coronal section showing vS2 lesion from an example animal, with S2 outline based on Allen Brain Atlas alignment via SHARP-track registration (Methods). **d,** Mean touch-evoked ΔF/F to whisker C2 protractions for all responsive neurons in an example mouse’s vS1 with neurons colored by touch type. Non-responsive neurons are excluded. Left, prior to any lesion. Middle, session after sham lesion. Right, session after vS2 lesion. **e,** Relative change in touch-evoked ΔF/F averaged across touch types in vS1 before and after vS2 lesion. Bar, mean (n=6). Grey, sham condition (n=9). P-values indicated for two-sided t-test of equality with 0 (dotted line): * p< 0.05, ** p < 0.01. **f,** As in **e** but broken up by touch type subpopulation.

To control for nonspecific effects, we preceded vS2 lesions with a ‘sham’ lesion of a non-vibrissal area (**Fig. 4b**). Following sham lesions, touch-evoked ΔF/F in vS1 remained unchanged (**Fig. 4e**, p=0.704). Sham lesions did not change the responsiveness of specific touch populations (**Fig. 4f;** USW, p=0.592; BSW, p=0.850; MW, p=0.550). We next asked whether vS2 lesions impacted vibrissal kinematics, as reduced touch intensity could account for the reduced response to touch. Touch count, the peak curvature during touch, and the peak velocity of whisking remained unchanged following vS2 lesions (**Extended Data Fig. 4**). We also did not find an overall change in correlation structure in vS1 after lesioning (**Extended Data Fig. 5**), suggesting that the pattern of touch-evoked activity in vS1 is not altered, but simply reduced in intensity. Finally, we looked at the whisker movement responsive cells (‘whisking’ neurons; Methods) and found that they do not decrease in number or responsiveness after lesions (**Extended Data Fig. 6**). Thus, the lesion effect is specific to the touch responsive population.

In sum, vS2 lesions result in a general decline in touch-evoked responsiveness in iso-somatotopic vS1. Nevertheless, neurons remain responsive to touch and the local correlation structure remains unaltered. Vibrissal S2 therefore non-selectively enhances the touch response in iso-somatotopic vS1.

### Vibrissal S1 lesions degrade vS2 touch responses in a whisker-specific manner

Vibrissal S2 enhances touch responses in somatotopically matched regions of vS1. If vS1 similarly enhances vS2 responses, this would imply that the two areas recurrently amplify spatially specific cortical touch responses. In contrast to vS2, vS1 somatotopy is sufficiently clear to target individual barrel columns^26^. We therefore recorded vS2 touch responses before and after focal lesions of a single vS1 barrel (**Fig. 5a-c**).

**Figure 5.**
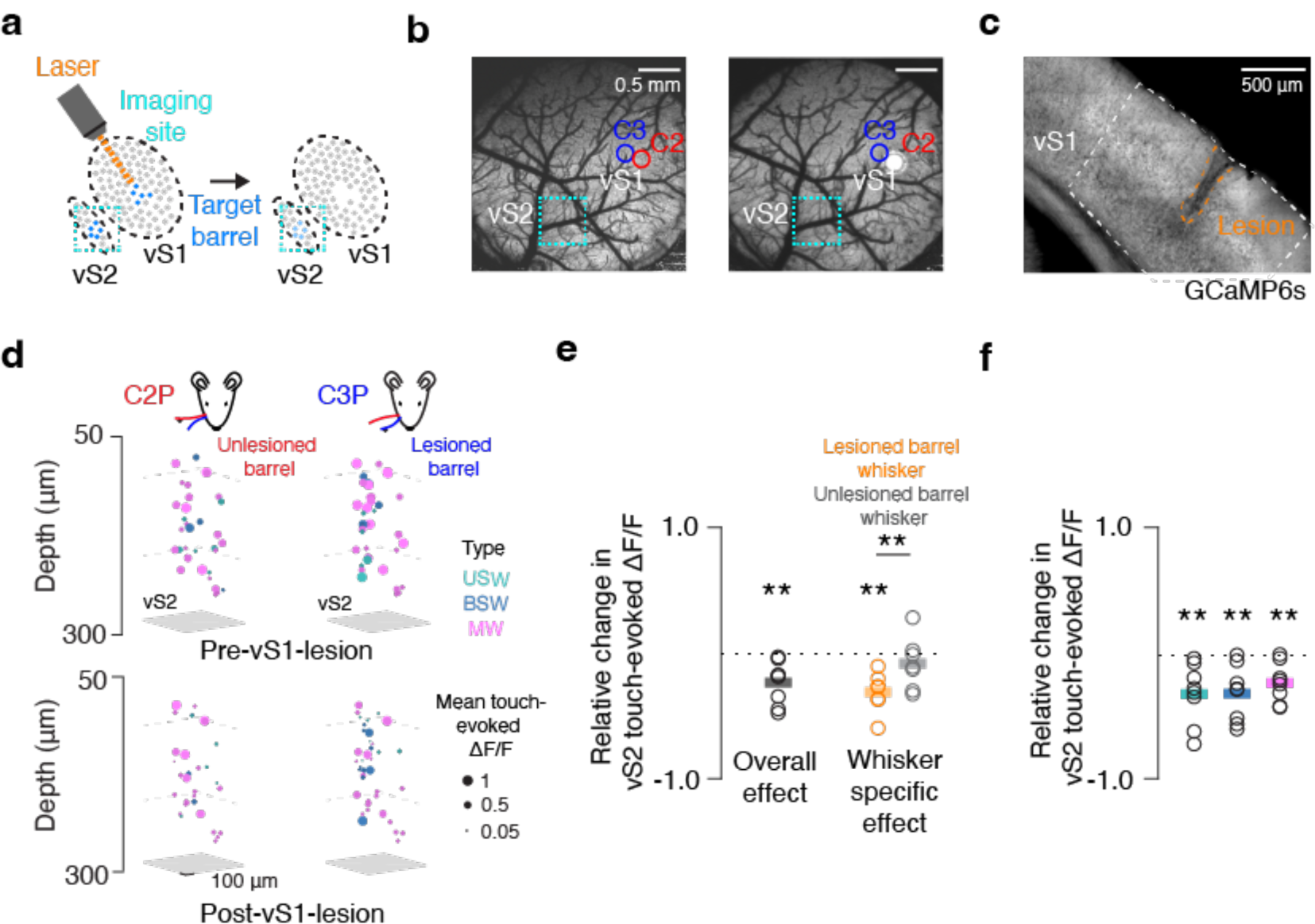
Touch response in vS2 declines after vS1 lesion. **a,** Lesion targeting single barrel in vS1. **b,** Widefield two-photon images of cranial window with barrel centers denoted before (left) and after (right) C2 barrel lesion. **c,** Coronal section showing vS1 lesion in an example animal, with vS1 outlined based on the Allen Brain Atlas. **d,** Mean touch-evoked ΔF/F for whisker C2 protractions (left) and whisker C3 protractions (right) for all responsive neurons in an example mouse with neurons colored by touch type in vS2 the session prior to (top) and after (bottom) a vS1 lesion targeting the whisker C3 barrel. **e,** Relative change in touch-evoked ΔF/F averaged across touch types in vS2 before and after vS1 lesion. Bar, mean (n=8). Orange, neurons that preferred to respond to touches by lesioned barrel’s preferred whisker. Grey, neurons that preferred the unlesioned barrel’s whisker. P-values indicated for two-sided paired t-tests: * p<0.05, **p<0.01. **f,** Same as in **e** but broken up by touch subpopulation.

We imaged a single 3-plane vS2 subvolume spanning 180 μm in total depth (**Extended Data Table 1**) in behaving mice both before and after lesion, yielding 1,244 ± 152 vS2 neurons in approximately 300-by-600 μm subregions across 8 mice (**Fig. 5d**). Lesioning a single whisker’s barrel column in vS1 reduced the relative mean touch-evoked ΔF/F in vS2 (**Fig. 5e**; p=0.007, t-test comparing relative change to 0). When we restricted our analysis to neurons preferring the whisker of either the lesioned or spared barrel, we found that the touch-evoked ΔF/F response to the lesioned barrel declined (**Fig. 5f**, p=0.002, n=7 mice), whereas the unlesioned barrel response did not (p=0.376, n=7 mice; lesioned vs. unlesioned, p=0.009). Reduced responses were observed in all three populations of touch neurons (USW, p=0.009; BSW, p=0.006; MW, p=0.006). Barrel-targeted vS1 lesions therefore produce a decline in aggregate vS2 touch responsiveness, with a larger effect among neurons responding to touch by the principal whisker of the targeted barrel. Despite this whisker specificity, the vS1 to vS2 projection is not selective among the classes of touch responsive neurons it influences.

In a subset of mice (n=7) with vS1 lesions, we evaluated the local lesion effect by imaging the adjacent tissue (1,610 ± 213 neurons). Though touch responses in the target barrel were eliminated, much of the touch response in the adjacent tissue remained, with a distance-dependent decline observed only for responses to the whisker of the targeted barrel (**Extended Data Fig. 7**). Traditional cortical lesions, which remove far larger volumes of tissue than our approach, can result in degradation of thalamocortical neurons^45,46^. We therefore examined several markers of thalamocortical degeneration: changes in Nissl stain reactivity, along with both microglial (Iba1) and astrocytic (GFAP) immunoreactivity (Methods). We did not observe any signs of thalamocortical degeneration (**Extended Data Fig. 8**). As with vS2 lesions, vS1 lesions did not alter vibrissal kinematics (**Extended Data Fig. 4**), the vS2 correlation structure (**Extended Data Fig. 5**), or the whisking responsive population (**Extended Data Fig. 6**). Reduced vS2 touch responsiveness following vS1 lesions is thus most likely due to the loss of direct input from vS1.

The vS1 to vS2 projection enhances the downstream touch response across touch cell types, with barrel-specific effects demonstrating the somatotopic specificity of this projection. Therefore, vS1-vS2 recurrently amplify touch responses across iso-somatotopic subregions of cortex.

## Discussion

We studied interactions across iso-somatotopic patches of L2/3 in mouse vS1 and vS2. We found that both vS1 and vS2 contain a sparse, mostly superficial, population of broadly tuned neurons that respond robustly to touch by both whiskers and exhibit high levels of synchrony (**Fig. 2**). This population carries a large proportion of the interareal touch signal in both directions, especially from vS2 to vS1, and its synchrony suggests that it is especially effective at influencing downstream targets^40^ (**Fig. 3**). Lesioning^44^ the subregion of either area that responded to touch by the spared whiskers resulted in a decline in touch response in the iso-somatotopic subregion of the other area, with whisker-specific vS1 lesions producing whisker-specific vS2 effects (**Figs. 4, 5**). We therefore propose that intracolumnar feedforward processing from L4 to L2 in vS1 and vS2 broadens receptive fields, yielding a sparse population of broadly tuned neurons that then recurrently amplify the touch response across iso-somatotopic patches of both areas.

Topography preserving excitatory projections between areas are common in cortex^4,47^. What computations might such circuits subserve? First, topography preserving circuitry could contribute to spatial attention^6,7^. Signals from higher order areas augmenting location specific responses in vS2, for instance, could propagate via such circuitry to produce enhanced responses in spatially matched regions of vS1 or other areas connected in this manner. Second, topography-respecting excitation could contribute to object recognition^9,10^ by elevating the sensitivity of topographically matched patches of cortex responding to the same stimulus. Because vS1 and vS2 encode distinct features of touch^27^, iso-somatotopic recurrent circuitry ensures that both responses concurrently reach higher order downstream areas putatively performing object recognition. Finally, such circuitry could act as a substrate to ‘bind’^8^ activity representing the same stimulus across disparate cortical regions, resulting in a cohesive percept.

We find that interareal projections are dominated by a sparse population of broadly tuned cells that presumably emerge due to feedforward processing from L4 to L2^37^. These cells are likely to be locally recurrently coupled^17^, resulting in elevated synchrony and greater downstream influence^40^. For the vS1 to vS2 projection, bidirectional single-whisker and multiwhisker cells together accounted for most projection activity, and for the vS2 to vS1 projection, multiwhisker cells constituted the outright majority of projection activity. Unidirectional single-whisker cells, though most numerous in both areas, contributed marginally. Our work suggests that iso-somatotopic recurrence across areas responding to specific touch stimuli is mediated by a broadly tuned population comprising 1-2% of L2/3 neurons in a given area^37^. This argues against interpretations of interareal connectivity in which feedforward projections are narrowly tuned and feedback projections are broadly tuned. This discrepancy could be due to actual differences between species and modalities, or it could be due to the failure to detect compare neurons in that are topographically matched across two areas to a sufficient degree, the oversampling of infragranular layers, and the failure to distinguish projecting neurons in many studies. Because we trim all but two of the ∼7 whiskers whose barrels our vS1 imaging window encompasses, some neurons that are multiwhisker are classified as single whisker, making it likely that the influence of multiwhisker cells is even more pronounced than reported here.

In visual cortex, feedback projections can suppress responses among iso-retinotopic patches of cortex^5,48,49^ and many interareal projections recruit inhibition in the target area^50,51^. Moreover, iso-retinotopic axonal projections appear to follow connectivity patterns that may preferentially amplify stimuli moving in specific ways across the visual field^52^. Particular pairs of areas may thus implement specific computations based on distinct patterns of both excitatory and inhibitory connectivity. Because of the difficulty of recording precisely matched sites across areas and the frequent lack of behavioral engagement and laminar specificity, it remains unclear if the discrepancy between our results and those observed in the visual system are due to genuine circuit differences or due to differences in experimental approach.

Though L2/3-to-L2/3 connectivity is observed among adjacent areas^19,22^, L5-to-L2/3, L2/3-to-L4, and other connections are also observed^3,4,20,53^. Modeling work suggests that projections connecting specific layers make distinct contributions to interareal interactions. Vibrissal S1 and vS2 are also interconnected across many layers: retrograde injections across the cortical depth of one area results in varying degrees of labeling in most layers of the other area^4^. Our retrograde injections were specifically targeted to L2/3, which resulted in most labeling being confined to L2/3 in the other area along with extensive intra-columnar labeling in other layers. This rich intra-columnar connectivity^54^ suggests that other layers indirectly contribute to the recurrent amplification revealed by our lesions. Moreover, both thalamocortical and corticothalamic projections are often topographically organized^55-57^, providing a further putative recurrent loop between vS1 and vS2. Finally, although our lesions typically target superficial layers^44^, dendrites of many neurons with somata below L2/3 will be cut and L2/3 input will be eliminated, so that nearly all neurons below the lesion site are likely to exhibit reduced touch responsiveness. It is thus likely that a substantial portion of the lesion effect is due to polysynaptic pathways, either within cortex or via thalamus. Projections between specific pairs of areas and laminae^53,58^ may contribute to distinct computations, with L2/3-mediated interactions driving recurrent amplification and other layer pairs potentially performing either complementary or distinct computations.

We find that vS1 and vS2 recurrently amplify the inter-areal cortical touch response largely via a sparse population of broadly tuned superficial L2/3 neurons that exhibit elevated synchrony and dominate the interareal projection touch response. During somatosensation, topography preserving projections thus augment the cortical response to localized touch across iso-somatotopic patches of vS1 and vS2.

### Methods Animals

Adult Ai162 (JAX 031562) X Slc17a7-Cre (JAX X 023527)^32^ mice (18 female, 21 male) were used (**Extended Data Table 1**). These mice express GCaMP6s exclusively in excitatory cortical neurons. Breeders were fed a diet that included doxycycline (625 mg/kg doxycycline; Teklad) so that mice received doxycycline until they were weaned, suppressing transgene expression throughout development. All animal procedures and protocols were approved by New York University’s University Animal Welfare Committee.

### Surgery

Cranial window and headbar implantation occurred in mice 6-10 weeks old under isoflurane anesthesia (3% induction, 1-2% maintenance). A dental drill (Midwest Tradition, FG 1/4’ drill bit) was used to make a circular craniotomy in the left hemisphere over vS1 and vS2 (3.5 mm in diameter; vS1 center: 3.5 mm lateral, 1.5 mm posterior from bregma; vS2 center: 4.7 mm lateral, 1.5 mm posterior from bregma). A triple-layer cranial window (4.5 mm external diameter, 3.5 mm inner diameter, #1.5 coverslip; two smaller windows were adhered to each other and the larger window with Norland 61 UV glue) was positioned over the craniotomy. The headbar and window were both affixed to the skull using dental acrylic (Orthojet, Lang Dental). Mice were post-operatively injected with 1 mg/kg of buprenorphine SR and 5 mg/kg of ketoprofen.

### Retrograde labeling

Retrograde viral injections were performed in untrained mice who had previously been implanted with a cranial window and had been trimmed to two whiskers (C2, C3). After surgical recovery, mice were run for one session on the imaging rig to identify either the touch patch in vS2 or the C2 and C3 barrels in vS1 (see Area Identification). For vS1 injections, we targeted the center of one of the two barrels (C2 or C3). For vS2 injections, we targeted the center of the patch of vS2 responsive to C2 and C3 touch. In both cases, the window was drilled off and a durotomy was performed. We injected 100 nL of pAAV-FLEX-tdTomato (Addgene, 28306-AAVrg, 1×10¹³ vg/mL diluted 1:50 in 1xPBS) into the target area at a depth of 200 μm and a rate of 20 nL/min (Narshige MO-10 hydraulic micromanipulator). Injection was performed using a glass capillary pulled with a micropipette puller (P-97, Sutter) and beveled to a tip with a ∼25**°** angle and 25 μm diameter. This was backfilled with mineral oil and 2 μL of the virus was pulled into the tip. The pipette was lowered into the target area at a rate of 300 μm/min which was followed by a 1-minute delay before injection began. An identical, new triple layer cranial window was placed over the craniotomy as before and re-affixed to the skull with dental acrylic. A subset of retrogradely injected animals were perfused (described below) and imaged on a confocal microscope (model SP5, Leica) using a 20x objective (**Extended Data Fig. 3**).

### Behavior

After surgical recovery, mice were water restricted and placed on a reverse light cycle. They were typically given 1 mL of water per day with small adjustments made to keep weight at 80-90% of pre-restriction baseline. Mice that had not been previously trimmed were trimmed to whiskers C2 and C3 and subsequently trimmed every 2-3 days.

Water-restricted mice were habituated to the behavioral apparatus for 2 days by head fixing them for 15-30 minutes and giving them free water. Mice were then trained on a two-whisker active touch task in which a pole was presented within range of one of two whiskers on every trial (**Fig. 1a**). Pole positions both in front of and behind each whisker’s natural resting position were used to encourage both retraction and protraction touches. The pole position on each trial was randomized but approximately half of trials in a given session targeted whisker C2 and the other half targeted whisker C3. Positions in front of and behind the whisker were used with as equal a frequency as possible. Pole positions were occasionally adjusted if animals changed their resting whisker position. For the longer whisker, pole placement was beyond the reach of the shorter whisker. Though the longer whisker did occasionally touch on trials meant for the shorter whisker, we obtained large numbers of isolated touch trials for all whiskers and touch directions (**Fig. 1d**). The pole was presented to the animal for one second, followed by a two second delay, and then a response cue (3.4 kHz, 50 ms) signaled the start of a one second response period. Mice would lick during the response period and would receive water randomly on ∼70% of trials. This was done to increase the number of trials during which the animal was engaged. A loud (60-70 dB) white noise sound was played for 50 ms following the onset of pole movement, which encouraged appropriately timed whisking. The lickport was moved along the anterior-posterior axis via a motor (Zaber) so that it was only accessible during the response period. Naïve mice whisked naturally at the onset of pole movement, likely due to the white noise sound, and would encounter the pole by chance. Over the course of a session, mice began to whisk vigorously in a stereotypical manner and subsequently lick after encountering the pole with the whisker.

A BPod state machine (Sanworks) and custom MATLAB software (MathWorks) running on a behavioral computer (System 76) controlled the behavioral task. Sounds were produced and controlled by an audio microcontroller (Bela). Three motorized actuators (Zaber) and an Arduino controlled lickport motion. Licks were detected via a custom detection circuit (Janelia).

### Whisker videography

Whisker video was acquired with custom MATLAB software using a CMOS camera (Ace-Python 500, Basler) with a telecentric lens (TitanTL, Edmund Optics) running at 400 Hz with 640 x 352 pixel frames. The video was illuminated by a pulsed 940 nm LED (SL162, Advanced Illumination) synchronized with the camera (typical exposure and illumination duration: 200 μs). 7-9 s of each trial were recorded, which included 1s prior to pole movement, the pole in-reach period, and several seconds following pole withdrawal. Data was processed on NYU’s High Performance Computing (HPC) cluster. Whiskers were detected using the Janelia Whisker Tracker^35^. Whisker identity assignment was then refined and evaluated using custom MATLAB software^17,30^. Whisker curvature (κ) and angle (θ) were then calculated at specific locations along the whisker’s length. Change in curvature, Δκ, was measured relative to a resting baseline curvature which was calculated at each angle independently. This value was obtained during periods when the pole was out of reach. Automatic touch detection was then performed, and touch assignment was curated manually using a custom MATLAB user interface^30^. Protractions were assigned negative Δκ values.

### Two-photon imaging

A custom MIMMS two-photon microscope (Janelia Research Campus; http://openwiki.janelia.org/ wiki/display/shareddesigns/MIMMS) with a 16X objective (Nikon) was used for cellular-resolution imaging. An 80 MHz titanium-sapphire femtosecond laser (Chameleon Ultra 2; Coherent) tuned to 940 nm was used, with powers out of the objective rarely exceeding 50 mW. The microscope included a Pockels cell (350-80-02, Conoptics), two galvanometer scanners (6SD11268, Cambridge Technology), a resonant scanner (6SC08KA040-02Y, Cambridge Technology), a 16x objective (N16XLWD-PF, Nikon), an emission filter for green fluorescence (FF01-510/84-30, Semrock), an emission filter for red fluorescence (FF01-650/60, Semrock), and two GaAsP PMTs (H10770PB-40, Hamamatsu) and two PMT shutters (VS14S1T1, Vincent Associates). A piezo (P-725KHDS; Physik Instrumente) was used for axial movement. Three imaging planes spanning 700-by-700 μm (512-by-512 pixels) and spaced at differential depths apart depending on the experiment were collected simultaneously at ∼7 Hz; we refer to this group of planes as a ‘subvolume’. Scanimage (version 2017; Vidrio Technologies) was used to collect all imaging data, and power was depth-adjusted in software using an exponential length constant of 250 μm. Up to 4 subvolumes were imaged per animal, and each subvolume was imaged for about 100 trials before moving to the next subvolume. All subvolumes were imaged on any given imaging day. After the first day of imaging, a motion-corrected mean image was created for each plane, which was then used as the reference image for any potential following imaging days. For animals where both vS1 and vS2 were imaged, two subvolumes were employed in each area spaced 60 μm apart, for a total span of 360 μm. For projection experiments, two subvolumes were collected in the non-injected area spaced 30-40 μm apart, for a total span of 180-240 μm. For lesion experiments, one subvolume was collected in each area spaced 60 μm apart, for a total span of 180 μm (**Extended Data Table 1**).

After acquisition, imaging data was processed on the NYU HPC cluster. First, image registration was completed for motion correction using a line-by-line registration algorithm^30^. Segmentation was performed on one session: neurons from the first day of imaging were detected using an automated algorithm based on template convolution that identified neuron centers, after which a neuron pixel assignment algorithm that detects annular ridges given a potential neuron center^31^ was used to identify the precise edges of the neuron. All pixels, including the nucleus, were used. This initial segmentation was manually curated, establishing a reference segmentation for each plane. On subsequent imaging days, the segmentation was algorithmically transferred to the new mean images for a given plane for that day^59^. After segmentation, ΔF/F computation and neuropil subtraction were performed. The neuropil-corrected ΔF/F trace was used for subsequent analyses.

In animals with projection labeling, neurons were classified as projecting if their mean tdTomato fluorescence exceeded a manually selected threshold. For each pixel on a plane, the cross-session mean tdTomato fluorescence was calculated. For any given neuron, its ‘redness’ was taken as the mean red fluorescence value for its constituent pixels in this mean image. A manual user interface that flagged neurons exceeding a threshold red fluorescence was used to find the threshold which appropriately partitioned tdTomato expressing neurons from non-expressing neurons in each animal.

### Lesions

A 1040 nm 80 MHz femtosecond laser (Fidelity HP, Coherent) was focused at a depth of 200–300 μm for 10–20 s at 1–1.5 W power (out of objective) to produce lesions. Vibrissal S1 lesions were made by centering the laser on a target barrel; vS2 lesions were made by centering the laser on the most responsive touch patch in the vS2 field of view. Sham lesions were performed in visual areas medial and posterior to vS1. Lesions were performed in awake, head fixed, durotomized animals sitting in the behavioral apparatus. Animals were monitored for signs of distress or discomfort. Typically, the lesion was performed at the end of a behavioral/imaging session. Post-lesion measurements were taken during the next session, approximately 24 hours after lesion. This timeline was chosen because it was determined that there was no significant recovery in the touch responses in the non-lesioned area if we imaged on timelines longer than 24 hours. This approach consistently yields lesions with a volume of 0.1-0.2 mm^3^, and when performed in animals expressing GCaMP6s, the radius of the post-lesion calcium response can be used to infer lesion extent^44^. In animals where intentionally large lesions were made (**Extended Data Fig. 8**), we made five or more additional lesions surrounding the initial lesion using the same depth, power, and timing parameters.

### Histology & Immunohistochemistry

After several days of imaging, some animals were perfused with paraformaldehyde (4% in PBS) and postfixed overnight. A vibratome (Leica) was used to cut coronal sections 100-mm thick which were mounted on glass slides with Vectashield antifade mounting media containing DAPI (Vector Laboratories). These sections were imaged on a fluorescent light microscope (VS120, Olympus). Slices were used to determine lesion location or injection spread. To ensure proper areal identification, all images were registered to the Allen Mouse Brain Common Coordinate framework using the SHARP-track pipeline^60^.

For **Extended Data Fig. 8**, two mice used in this study were perfused 72 hours after vS1 lesion (after all imaging was complete). Two additional mice were given exorbitantly large vS1 lesions (see: Lesions) and perfused 72 hours after lesion. A vibratome (Leica) was used to cut 50-μm thick sections and sections that included the lesion were either used for immunohistochemistry or Nissl staining.

For immunohistochemistry, slices were incubated overnight under agitation with primary antibody that was made in 1% bovine serum albumin and 0.05% sodium azide. Slices were labeled with either rabbit anti-Iba1 (Sigma; NC9288364) or mouse anti-GFAP (glial fibrillary acidic protein; Sigma G3893)^61^ antibodies. After incubation, slices were washed and incubated in secondary antibody (1:500) conjugated to Alexa Fluor 647 (Sigma). Finally, slices were rinsed and mounted using an antifade mounting media (Vector Laboratories), and subsequently imaged using an Olympus VS120 microscope and a Leica SP5 confocal microscope. Primary antibodies: rabbit anti-Iba1 (019-19741; Wako; 1:500 dilution), mouse monoclonal anti-GFAP (G3893; Sigma-Aldrich; 1:1,000 dilution). Secondary antibodies: goat anti-Mouse, Alexa Fluor 647 (Iba1; Thermo Fisher A-21244), and Goat anti-Rabbit, Alexa Fluor 647 (GFAP; Thermo Fisher A-21235).

For Nissl staining, slices were immediately mounted and then deparaffinized with xylene, and sequentially rinsed in varying concentrations of ethanol and distilled water. Sections were then Nissl stained by submersion of slides in 0.125% cresyl violet, followed by further rinsing. Dried and stained slides were imaged on the Olympus VS120 microscope in bright-field mode.

### Area identification

The locations of vS1 (including individual barrel locations) and vS2 were identified by measuring the GCaMP6s ΔF/F at coarse resolution (4x objective, Nikon; field of view, 2.2 x 2.2 mm) on the two-photon microscope while the whiskers were deflected individually with a pole. Imaging was performed for a single imaging plane at 28 Hz. This was done in awake mice not engaged in any task. For injections and lesions, we then briefly imaged at cellular resolution using volumetric imaging to further restrict ourselves to the relevant injection or lesion target. Whiskers were individually deflected with the pole for approximately 100-200 trials while we imaged in one subvolume with three planes and 60 μm spacing. We then analyzed the touch response in individual neurons (as described below), identifying a single barrel for vS1 injections or lesions, or the vS2 touch patch.

Vibrissal S1 and vS2 are similarly extensive along the anterior-posterior (AP) axis, but vS2 is relatively compressed along the mediolateral (ML) axis^4,25^. Within our imaging fields of view we therefore selected a 600-by-600 μm field of view for vS1 centered on the C2/C3 border, and a 600 μm (AP axis) by 300 μm (ML axis) field of view for vS2. This ensured that we were not analyzing vS1 cells in our vS2 field of view and that we did not underestimate vS2 cell fractions. This restriction included the area in vS2 with retrogradely labeled neurons following vS1 injections.

### Touch and whisking responsiveness

A neuron was classified as responsive or non-responsive for a particular touch trial by comparing ΔF/F_baseline_, the mean ΔF/F for the 6 frames (0.85 s) preceding the first touch, to ΔF/F_post-touch_, the mean ΔF/F for the period between the first touch and two frames after the final (interframe interval, ∼143 ms). For each neuron, we computed the standard deviation of ΔF/F across all pre-touch frames (6 frames prior to first touch), yielding a noise estimate, σ_baseline_. A neuron was considered responsive on a trial if the ΔF/F_post-touch_ exceeded ΔF/F_baseline_ for that trial by at least 2*σ_baseline_. Neurons that were responsive on at least 5% of trials for a given touch type were considered part of the responsive pool for that type.

To compute a neuron’s mean touch-evoked ΔF/F, we incorporated the response to all touch types for which that neuron was significantly responsive. In all cases, touch-evoked ΔF/F was measured as ΔF/F_post-touch_-ΔF/F_baseline_ for a given trial. For a given touch type (C2P, C2R, C3P, and C3R), mean touch-evoked ΔF/F was calculated using trials where only that touch type occurred. For unidirectional single whisker cells, we used the mean touch-evoked ΔF/F for the single touch type it responded to. For bidirectional single whisker cells, we used the mean touch-evoked ΔF/F across the two directions for the whisker the cell responded to, weighing both touch types equally. Finally, for multiwhisker cells, we first calculated the mean touch-evoked ΔF/F for the touch types to which that cell responded, then took the mean across these values.

Whisking responsive neurons (**Extended Data Fig. 6**) were classified in the same way as described for touch responsive cells, with the key difference of aligning responses based on whisking bout onset as opposed to touch. Whisking bouts were defined as periods where the amplitude of whisking derived from the Hilbert transform exceeded 5°. As before, ΔF/F_baseline_ and ΔF/F_post-whisking-onset_ were compared and a cell was considered whisker responsive overall if it was responsive on at least 5% of trials with whisking.

### Correlation analysis

Pearson correlations were calculated across neuron pairs either around the time of touch or restricted to time points outside of touch (‘spontaneous’ epoch). In all cases, correlations were computed using the ΔF/F values at all included timepoints. Periods outside of touch were defined as any timepoints that did not fall between 1 s before and 10 s after a touch. For touch correlations, a mean correlation was calculated using a window that began 1 s prior to first touch onset and ending 4 s after final touch offset. Four correlations were computer per neuron, one each for C2P, C2R, C3P, and C3R touches. For unidirectional single-whisker neurons, we only used the value for the touch type the neuron responded to. For bidirectional single-whisker and multiwhisker neurons, we took the grand mean of the mean correlations from trials with isolated touches for the touch types the neuron was considered responsive to.

Because only individual subvolumes were recorded simultaneously, we aggregated across subvolumes in some cases. For non-projection analyses, we only employed the superficial subvolume. For projections, we imaged superficial L2/3 with two subvolumes due to the relatively low yield of retrograde labeling, and so we aggregated these when possible within animals by using the grand mean of individual population correlations across subvolumes. We applied a 5 neuron minimum within a given category for correlation computation. This did not impact most analyses, but for projection-based correlations, it precluded the analysis of bidirectional single-whisker neurons in vS2 altogether and did result in the exclusion of some subvolumes. If only one subvolume met this criterion for an animal, that subvolume was used as the correlation for that animal. If both subvolumes met the criteria, correlations for that animal were computed as the mean across the two subvolumes. If both subvolumes failed to meet the cell count criteria, the animal was excluded from analysis.

### Analysis of the fraction of response carried by each population

For the analyses in **Figs. 2g, 2h** and **3g**, for each animal, we found the response of each touch neuron (**Fig. 2**) or each projecting neuron (**Fig. 3**) to each type of touch that neuron responded to for every touch trial. For each trial, we took the mean response to touch for each type of touch that the neuron responded to, and that mean was the mean response to touch for that neuron on a given trial. For each neuron we took the mean across trials, and then sorted neurons based on their touch response classification (USW, BSW, or MW). By summing the mean response of each neuron in each group we found the overall mean response to touch in each touch type category. We took the sum of the mean response of all touch or projecting neurons to be the total touch response of the touch or projecting neurons. The division of the mean response to touch in each category with the total response in an area gives the fraction of response carried by each cell type in that area.

## Statistical analysis

We used the two-tailed paired t-test for comparisons across matched groups, where pairing was typically within-animal. The two-sample t-test was used to compare between distinct samples. In a few cases, we used a t-test against 0 or 1 to compare population change to chance. All statistical analysis was performed using MATLAB, and a list of exactly which animals are used in each experiment can be found in **Extended Data Table 1**.

## Data availability

All data from this study will be made available on our laboratory website upon publication.

## Code availability

Code used to generate the figures in this study will be made available via the laboratory github page upon publication.

## Acknowledgements

We thank Alisha Ahmed, Andy Garcia, Ravi Pancholi and Leopoldo Petreanu for comments on the manuscript. We thank Rob Froemke, Anthony Movshon, Dan Sanes and David Schneider for discussion. This work was supported by the Whitehall Foundation and the National Institutes of Health (R01NS117536).

## Author Contributions

L.R. and S.P. designed the study. L.R. carried out most experiments with help from A.S. and M.L. L.R. performed data analysis. L.R. and S.P. and wrote the paper.

## Supplementary Materials

**Extended Data Table 1.** Animal list. All mice were transgenic adult Ai162 X Slc17a7-Cre and therefore expressed GCaMP6s only in excitatory neurons^32^. Spared whiskers were C2/C3 for all animals. We used a mix of male and female animals. Age of imaging onset is provided in weeks. Imaging scheme indicates the net depth span imaged across 1 or 2 subvolumes (‘sv.’). Cell counts represent all neurons imaged per area and do not factor in restricted subregions (Methods). In general, each figure represents one key experiment and animals listed as part of that experiment comprise the dataset for each figure: vS1-vS2 comparison with broad depth sampling, ‘vS1 vs. vS2’ (Fig. 2); lesion experiments (**Figs. 3, 4**). For retrograde injection animals (‘retro. inj’; Fig. 3), the injected area is specified; imaging took place in the other area. ‘Rad. eff.’ indicates animals included in the radius of effect analysis (**Extended Data Figure 7**).

| Mouse | Sex | Age (wk) | Areas imaged | Imaging depth span | Neuron count | Experiments |
| --- | --- | --- | --- | --- | --- | --- |
| 14351 | F | 12 | vS1/vS2 | 360 μm (2 sv.) | vS1: 4,416<br>vS2: 3,567 | vS1 vs. vS2, vS1 lesion |
| 14359 | F | 16 | vS1/vS2 | 360 μm (2 sv.) | vS1: 4,578<br>vS2: 3,509 | vS1 vs. vS2, vS1 lesion, vS1 retro. inj. |
| 17722 | F | 12 | vS1/vS2 | 180 μm (1 sv.) | vS1: 2,865<br>vS2: 2,082 | vS1 lesion, rad. eff. |
| 18921 | M | 11 | vS1/vS2 | 180 μm (1 sv.) | vS1: 3,277<br>vS2: 2,509 | vS1 lesion, rad. eff. |
| 14332 | F | 14 | vS1/vS2 | 180 μm (1 sv.) | vS1: 3,454<br>vS2: 1,741 | vS1 lesion, rad. eff. |
| 18489 | F | 16 | vS1/vS2 | 180 μm (1 sv.) | vS1: 2,493<br>vS2: 2,072 | vS1 lesion, rad. eff. |
| 18920 | M | 16 | vS1/vS2 | 180 μm (1 sv.) | vS1: 2,904<br>vS2: 2,017 | vS1 lesion, rad. eff. |
| 20035 | F | 18 | vS2 | 180 μm (2 sv.) | vS2: 4,138 | vS1 retro. inj. |
| 20428 | M | 14 | vS2 | 240 μm (2 sv.) | vS2: 4,092 | vS1 retro. inj. |
| 20047 | F | 12 | vS2 | 180 μm (2 sv.) | vS2: 3,157 | vS1 retro. inj. |
| 20044 | M | 12 | vS2 | 240 μm (2 sv.) | vS2: 3,088 | vS1 retro. inj. |
| 19806 | F | 13 | vS2 | 180 μm (2 sv.) | vS2: 2,940 | vS1 retro. inj. |
| 19821 | F | 23 | vS2 | 180 μm (2 sv.) | vS2: 3,954 | vS1 retro. inj. |
| 16623 | F | 16 | vS1/vS2 | 360 μm (2 sv.) | vS1: 4,868<br>vS2: 3,426 | vS1 vs. vS2, vS2 lesion, sham lesion |
| 14362 | F | 16 | vS1 | 360 μm (2 sv.) | vS1: 5,159 | vS2 retro. inj. |
| 17517 | M | 17 | vS1/vS2 | 360 μm (2 sv.) | vS1: 4,128<br>vS2: 3,301 | vS1 vs. vS2, sham lesion |
| 17721 | F | 21 | vS1/vS2 | 180 μm (1 sv.) | vS1: 2,511<br>vS2: 1,440 | vS2 lesion, sham lesion |
| 18927 | M | 13 | vS1/vS2 | 180 μm (1 sv.) | vS1: 2,676<br>vS2: 2,429 | vS2 lesion, sham lesion |
| 18933 | M | 13 | vS1/vS2 | 180 μm (1 sv.) | vS1: 3,080<br>vS2: 1,905 | vS2 lesion |
| 20048 | F | 17 | vS1/vS2 | 180 μm (1 sv.) | vS1: 2,720<br>vS2: 1,939 | vS2 lesion |
| 18922 | M | 17 | vS1/vS2 | 180 μm (1 sv.) | vS1: 2,843<br>vS2: 2,205 | vS2 lesion |
| 18912 | M | 16 | vS1 | 180 μm (2 sv.) | vS1: 5,136 | vS2 retro. inj. |
| 18919 | M | 16 | vS1 | 240 μm (2 sv.) | vS1: 3,456 | vS2 retro. inj. |
| 20055 | F | 16 | vS1 | 180 μm (2 sv.) | vS1: 5,123 | vS2 retro. inj. |
| 19781 | M | 18 | vS1 | 240 μm (2 sv.) | vS1: 5,612 | Sham lesion, vS2 retro. inj. |
| 19770 | M | 15 | vS1 | 180 μm (2 sv.) | vS1: 5,315 | Sham lesion, vS2 retro. inj. |
| 20421 | M | 13 | vS1 | 180 μm (2 sv.) | vS1: 4,697 | vS2 retro. inj. |
| 20430 | M | 16 | vS1 | 180 μm (2 sv.) | vS1: 3,559 | vS2 retro. inj. |
| 19776 | M | 23 | vS1 | 240 μm (2 sv.) | vS1: 6,254 | vS2 retro. inj. |
| 16650 | M | 10 | vS1/vS2 | 360 μm (2 sv.) | vS1: 5,191<br>vS2: 5,163 | vS1 vs. vS2 |
| 16652 | M | 12 | vS1/vS2 | 360 μm (2 sv.) | vS1: 3,146<br>vS2: 4,045 | vS1 vs. vS2 |
| 14363 | M | 17 | vS1/vS2 | 360 μm (2 sv.) | vS1: 4,141<br>vS2: 3,649 | vS1 vs. vS2 |
| 17510 | M | 18 | vS1/vS2 | 360 μm (2 sv.) | vS1: 5,644<br>vS2: 3,047 | vS1 vs. vS2 |
| 17518 | M | 17 | vS1/vS2 | 360 μm (2 sv.) | vS1: 5,160<br>vS2: 2,616 | vS1 vs. vS2 |
| 14353 | M | 12 | vS1/vS2 | 180 μm (1 sv.) | vS1: 2,804<br>vS2: 2,293 | Sham lesion |
| 14355 | F | 13 | vS1/vS2 | 180 μm (1 sv.) | vS1: 2,315<br>vS2: 2,193 | Rad. eff., vS1 retro. inj. |
| 14354 | F | 13 | vS1/vS2 | 180 μm (1 sv.) | vS1: 2,036<br>vS2: 1,467 | Sham lesion, vS1 retro. inj. |
| 14335 | F | 12 | vS1/vS2 | 180 μm (1 sv.) | vS1: 2,839<br>vS2: 1,406 | Sham lesion |
| 18490 | F | 13 | vS1/vS2 | 180 μm (1 sv.) | vS1: 3,032<br>vS2: 2,039 | vS1 lesion, rad. eff. |

**Extended Data Figure 1.**
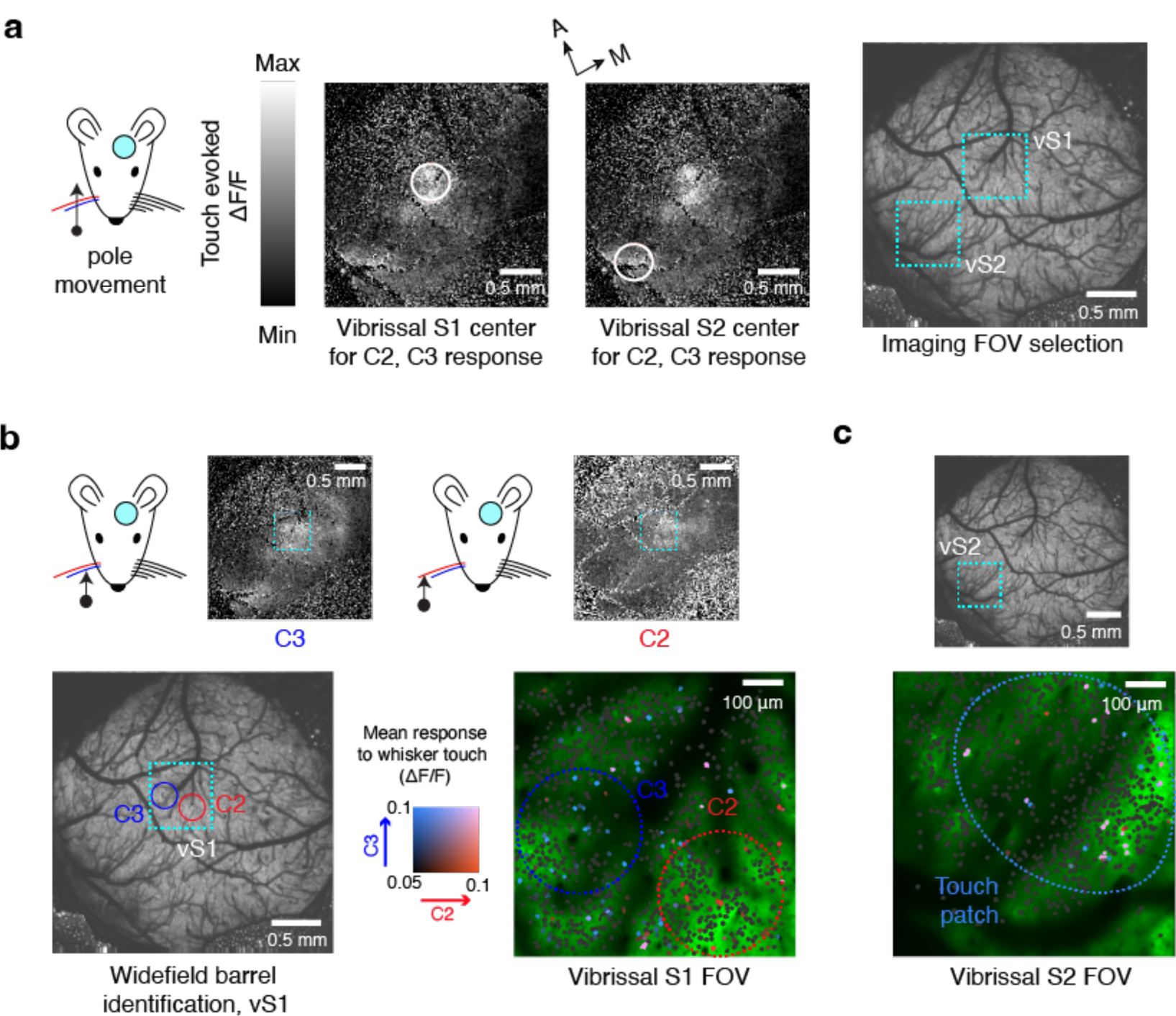
Vibrissal S1 barrel and vS2 touch patch identification. **a,** Left to right; cartoon showing coarse stimulation of both spared whiskers at once via a pole pushing against the two whiskers; touch evoked widefield calcium response to such stimulation, with manually selected centers of vS1 and vS2 delimited in white; widefield 2-photon image of a cranial window with vS1 and vS2 fields of view outlined in cyan. **b,** Top, same as in **a** but for stimulation of individual whiskers, revealing C3 and C2 barrel centers, again manually selected. Bottom, left to right: barrels within widefield image and vS1 imaging FOV; touch map with neurons colored by whisker preference with these barrels overlaid. **c,** Cellular-resolution touch-map for vS2 field of view in the same animal, with vS2 touch patch overlaid.

**Extended Data Figure 2.**
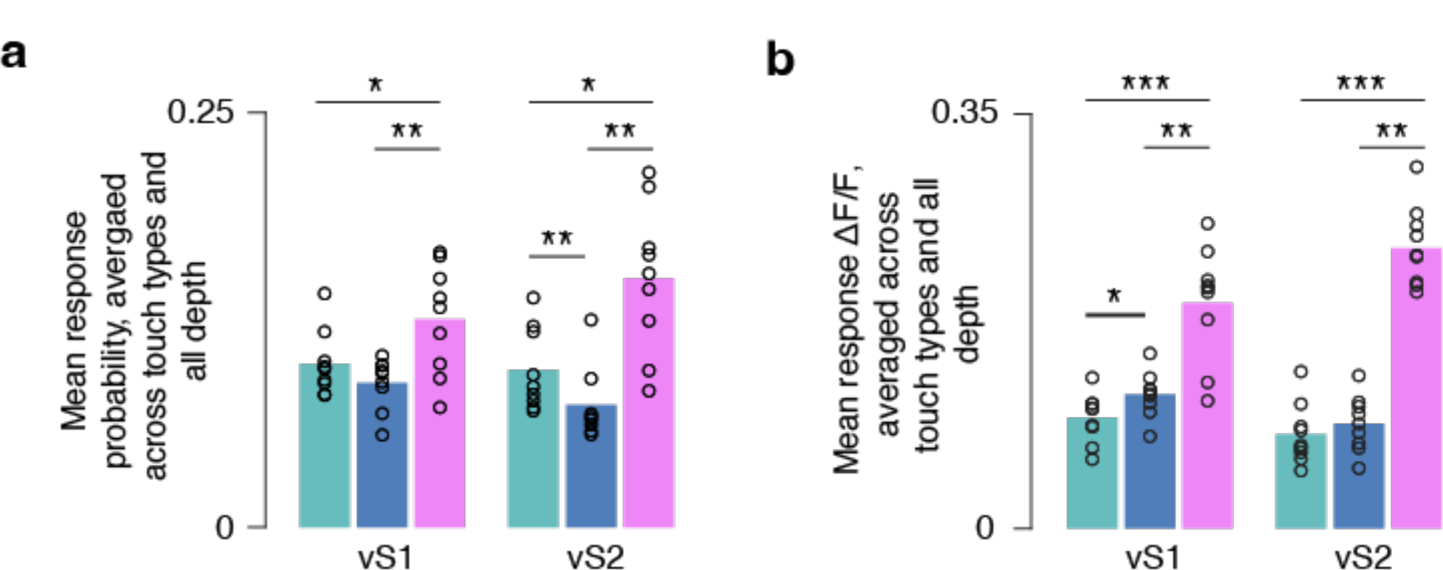
Contribution of specific touch cell types to touch response across all depths. **a,** Mean response probability for each type in each area averaged across all touch types for which individual neurons were deemed responsive, including all sub-selected neurons across all depth (not restricted to superficial subvolumes as in Fig. 2e). **b,** Mean touch-evoked ΔF/F averaged across touch types for which individual neurons were deemed responsive, across all depths (not superficially restricted like Fig. 2f). P-values indicated for paired t-test: * p < 0.05; ** p < 0.01; *** p < 0.001.

**Extended Data Figure 3.**
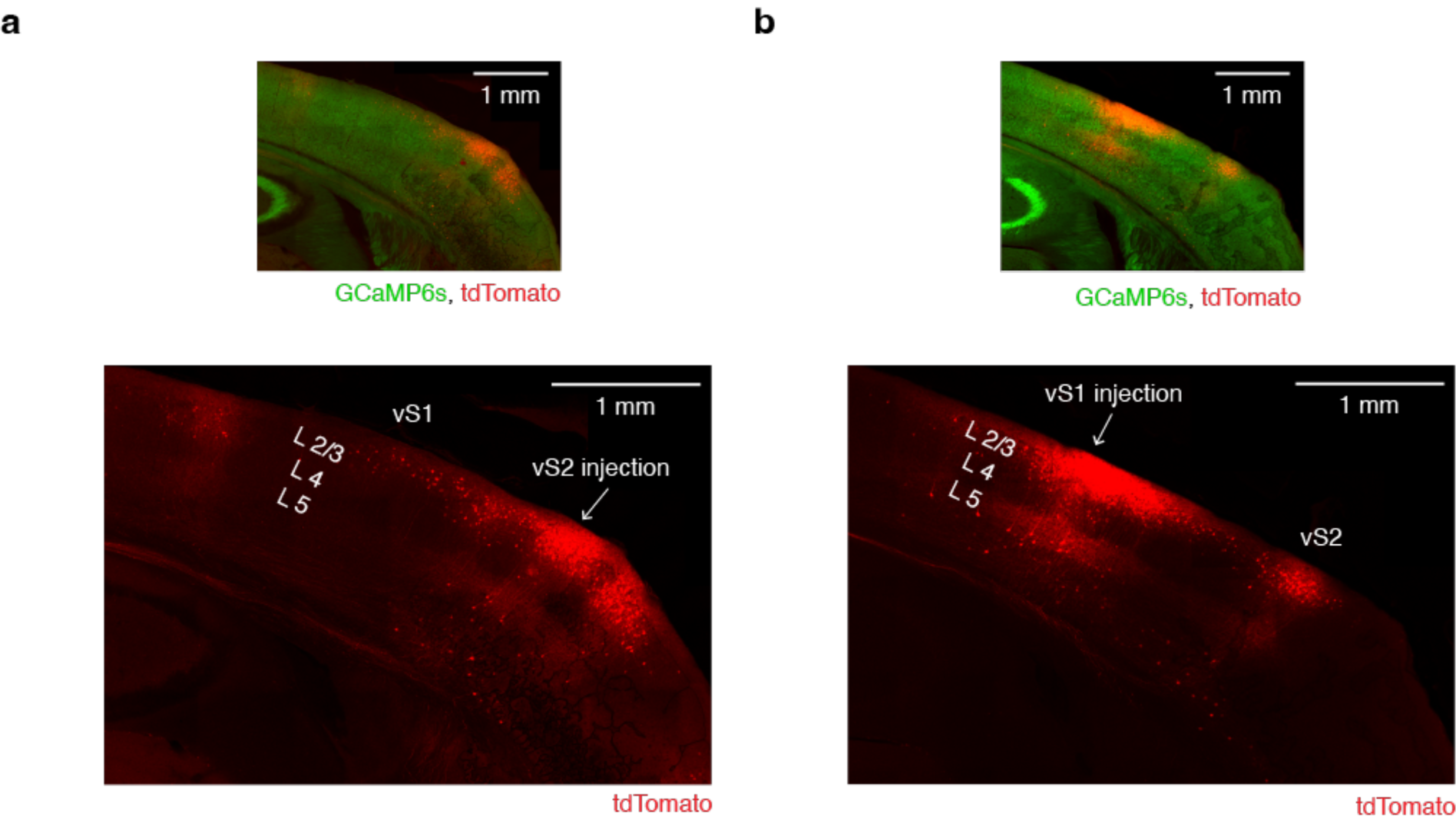
Distribution of retrogradely labeled neurons following viral injection into either vS2 or vS1. **a,** Top: overlay of red and green channels from a confocal image displaying a retrograde injection in L2/3 of vS2 (Methods). Bottom: Red channel separated, 20x zoom. Injection site denoted. **b,** Same but for an injection in L2/3 of vS1.

**Extended Data Figure 4.**
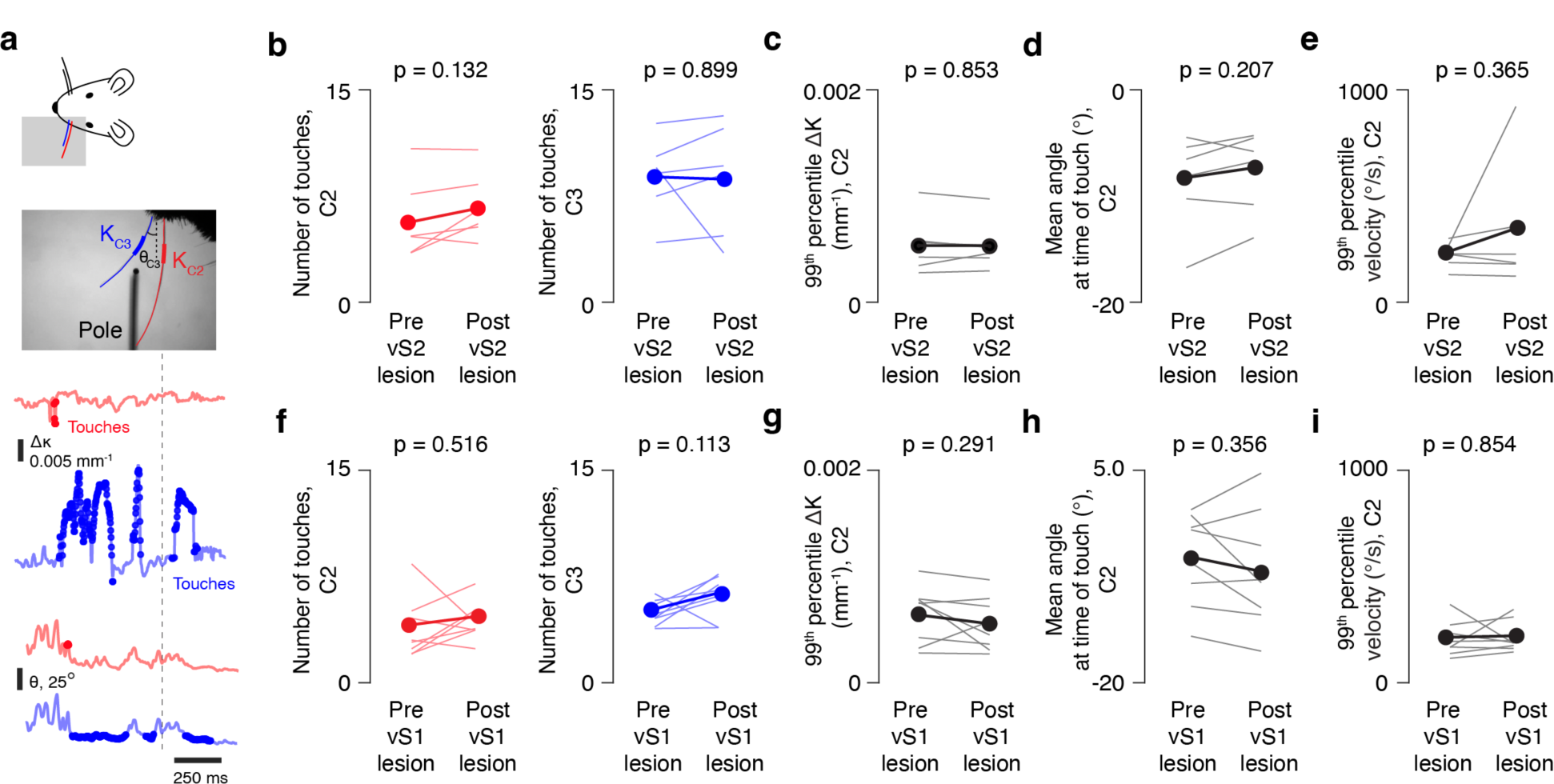
Lesions of vS1 and vS2 do not impact vibrissal kinematics. **a,** Kinematic variables. Top, example whisker video frame, showing curvature (Δκ) measurement for both whiskers and angle (θ) measurement for C3. Bottom: Δκ and θ traces for both whiskers with touches overlaid. **b,** Left (red), mean number of touches made by C2 across trials before and after vS2 lesions. Thick line: cross animal mean. Thin lines: individual mice, n=6. P-values indicated for two-sided paired t-test. Right (blue), same but for C3. **c,** Peak change in curvature (Δκ) of C2 before and after vS2 lesion. ‘Peak’ is calculated by finding the 99^th^ percentile of values across all whisker contact epochs. **d,** Mean angle of C2 at the time of whisker touch before and after vS2 lesion, averaged across all times during which the pole was touched. **e,** Peak velocity of C2 before and after vS2 lesion. Velocity is computed by comparing two subsequent frames (Methods), and ‘peak’ is calculated as the 99^th^ percentile of values when the pole is in reach but prior to the first touch. **f-i**, same as in **b-e** but comparing kinematics before and after vS1 lesions, n=8 mice.

**Extended Data Figure 5.**
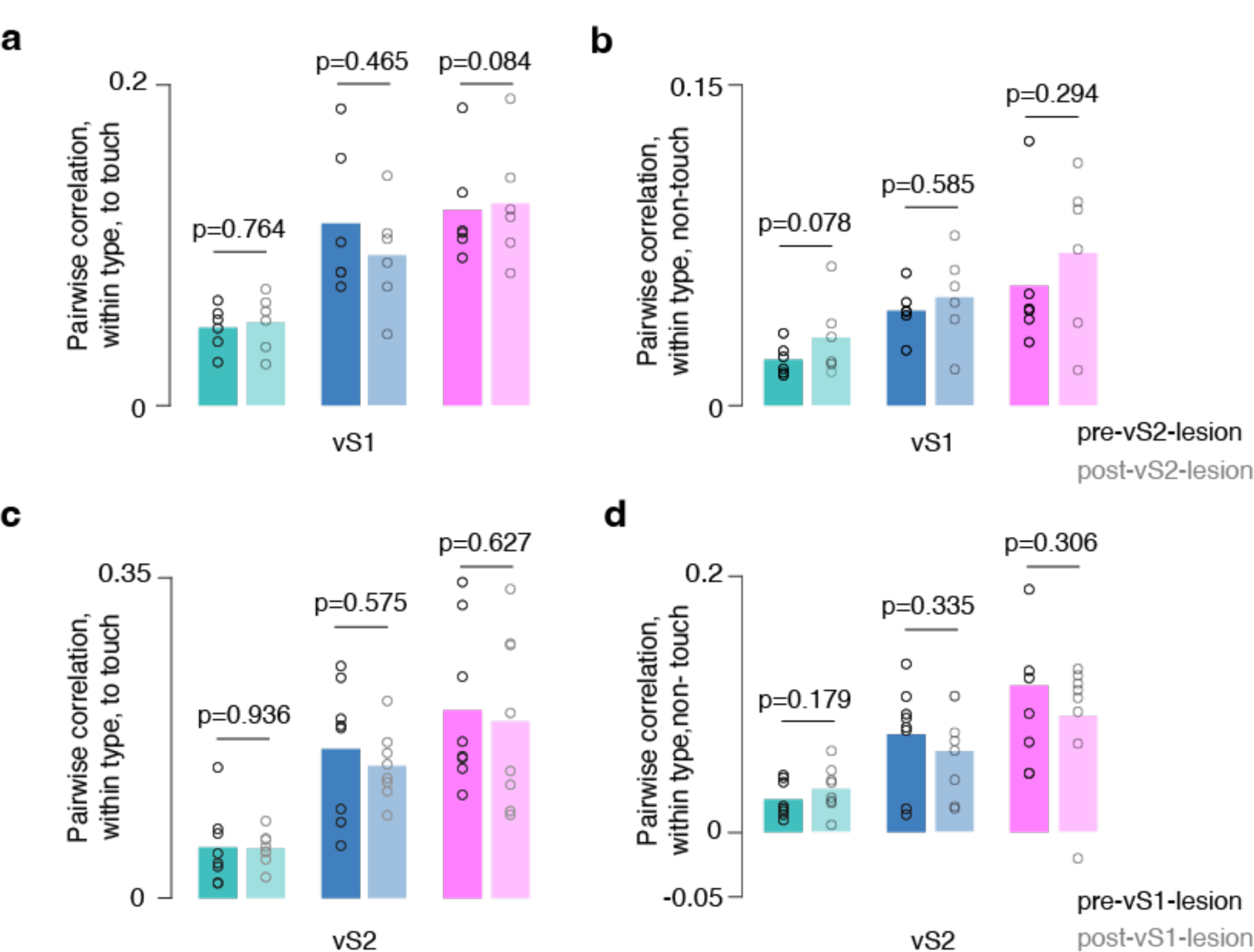
Columnar-scale lesions of vS1 and vS2 do not lead to overall changes in the correlation structure in the other area. **a,** Mean within-type pairwise correlations in vS1 during the period around touch by group, pre and post vS2 lesion. Bars: cross-animal mean, circles: individual animal means. P-values are for two-sided paired t-test comparing mean correlation before and after vS2 lesions, n=6 mice. **b,** Mean within-type pairwise ‘spontaneous’ correlations during non-touch period in vS1 before and after vS2 lesion. **c,** Same as **a** but for vS2 before and after vS1 single barrel lesions; n=8 mice. **d**, Same as in **b** for vS2 before and after vS1 lesion.

**Extended Data Figure 6.**
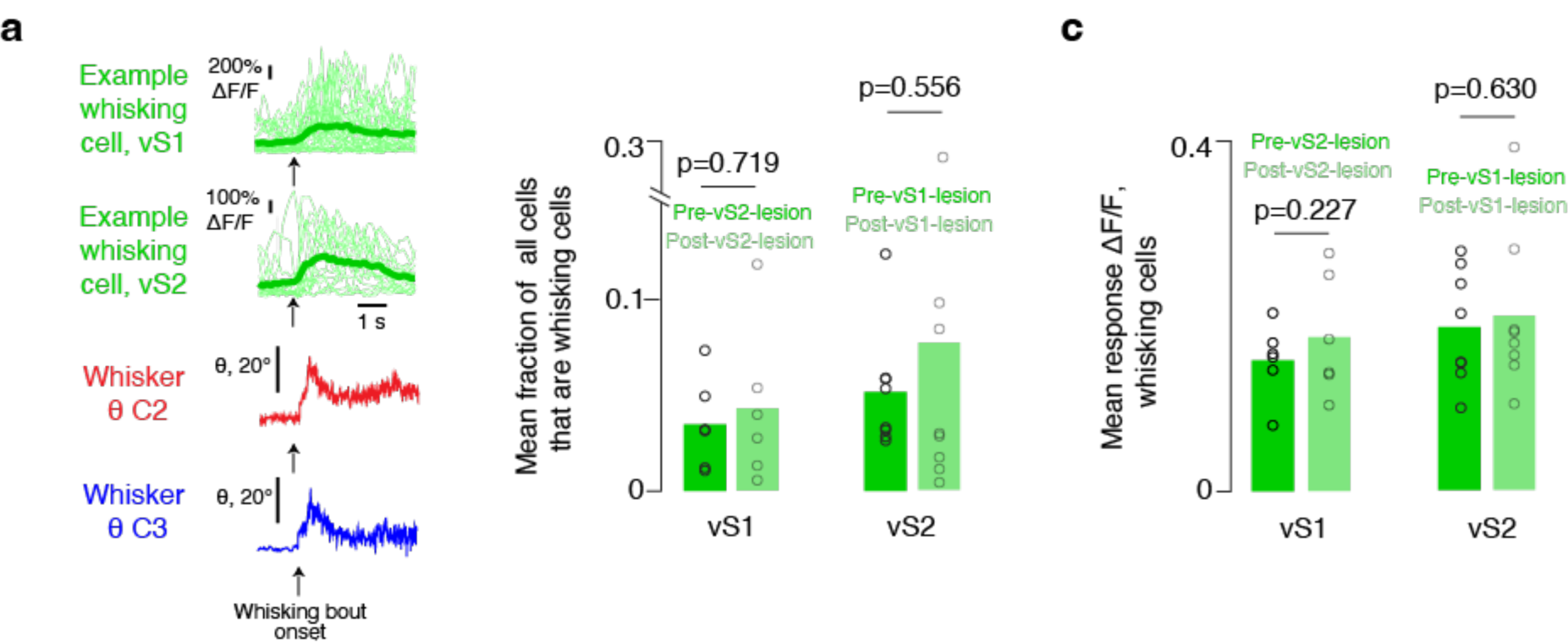
Columnar-scale lesions of vS1 and vS2 do not alter the whisking population in the other area. **a,** ΔF/F traces for two example whisking cells, one from vS1 and one from vS2. Thin lines, individual trials; thick lines, mean. Mean angle traces for C2 and C3 shown below, all aligned to whisking bout onset. **b**, Mean fraction of cells in each area that are responsive to whisking (Methods), before and after lesion. **c,** Mean response ΔF/F of whisking cells before and after lesion in both areas.

**Extended Data Figure 7.**
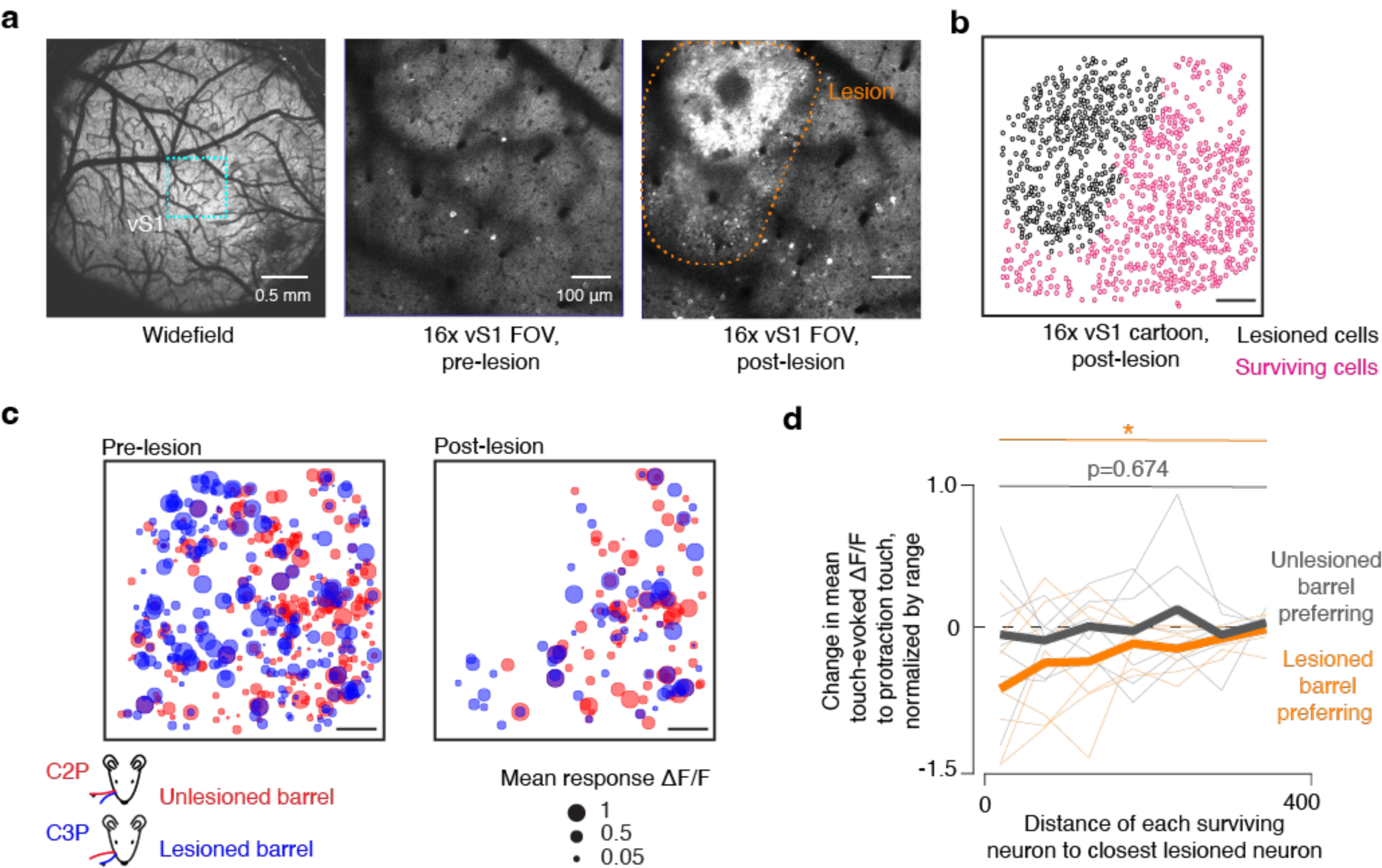
Radius of effect of single vS1 barrel lesion. **a,** Left, vS1 field of view shown in widefield 2-photon image. Middle, vS1 imaging field of view at 16x. Right, same 16x field of view but after lesion to one barrel. **b,** Cartoon of the same FOV showing which cells we determined to be dead after lesion (black) versus cells that survived. **c,** Map of touch-evoked ΔF/F for touch responsive neurons in vS1 to whisker C2 protraction (red) and whisker C3 protraction (blue) before (left) and after (right) lesion of the C3 barrel in vS1. **d,** Normalized change in the mean protraction-touch-evoked ΔF/F as a function of each surviving neuron’s distance to the closest lesioned neuron. Distance bins are 50 μm. Thin lines, mean change across neurons in a given distance bin for a single animal. Thick lines, cross animal mean. Orange, neurons responded most strongly to whisker touch of the lesioned barrel’s whisker. Grey, neurons that preferred to respond to the unlesioned barrel’s whisker. P-values indicated for paired t-tests comparing distance bin 1 to the final distance bin for unlesioned and lesioned whisker responsive neurons; * p <0.05.

**Extended Data Figure 8.**
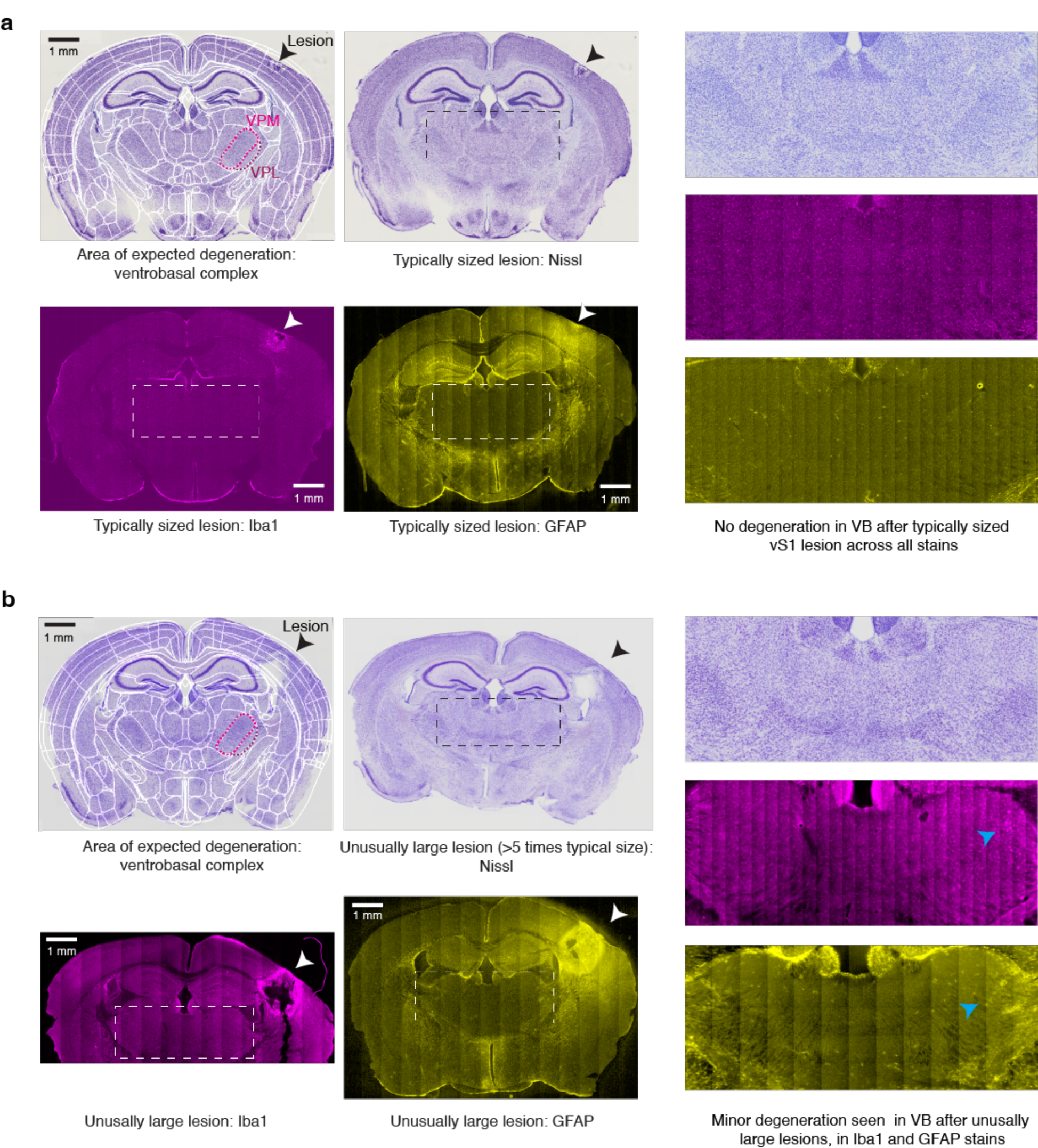
Impact of columnar-scale vS1 lesions on thalamus. **a,** Immunohistochemistry performed a few days after lesioning in example animals with normally sized lesions of vS1. Top left, Nissl stain, first overlaid with anatomical mapping from the Allen Brain Atlas, with thalamic areas in which we might expect degeneration (VB, including VPM and VL), outlined. Arrow shows lesion in vS1. Bottom left: Iba1 stained slice (magenta) and GFAP stained slice (yellow) with lesion shown. Right: zoomed in view of all three slices, corresponding to outlined box on the left. **b,** Same as in a but for example animals (not used for imaging or lesion experiments) in which we made intentionally large lesions, larger than 5x the size of our normal sized lesions. Many of these lesions also went through the white matter. Blue arrows denote possible thalamic degeneration.

